# Fluorescent Biosensor-Guided Engineering of Enzyme Cascades for Electrochemical Applications

**DOI:** 10.1101/2025.11.21.689824

**Authors:** Nathan J. Ricks, Monica Brachi, FNU Arun, Egor Baiarashov, Shelley D. Minteer, Ming C. Hammond

## Abstract

Substrate channeling is a strategy for enhancing flux and yield in enzymatic cascades and is increasingly relevant for applications in biocatalysis, biotechnology, and bioelectrochemical systems. However, efforts to engineer channeling are limited by the lack of high-throughput methods to evaluate and optimize channeling efficiency. Here, we present a fluorescence-based screening assay to rapidly assess substrate channeling in a model system involving fumarase and malate dehydrogenase, two sequential enzymes from the Krebs cycle. By expressing genetic fusions in *E. coli*, quantifying intermediate (malate) and product (NADH) formation in lysate using orthogonal fluorescent readouts, and comparing product-to-intermediate ratios, we screened a library of linker variants designed to promote electrostatic channeling. A top-performing construct was identified and validated through classical channeling assays. This hit demonstrated increased product yield and current output when immobilized on electrodes with a bilayer architecture, highlighting utility in bioelectrocatalysis. We further showed that the channeling linker could be applied to a *de novo* designed single-chain fumarase, which preserved channeling capability and exhibited improved thermal stability. These results establish a generalizable and scalable method for engineering and evolving substrate channeling, with broad implications for pathway optimization and enzyme design in synthetic biology, bioprocessing, and energy applications.

## INTRODUCTION

Redox enzymes are essential in many metabolic cascades including biosynthesis, energy conversion, and nutrient metabolism. Given their efficiency, these enzymes have been explored as electrocatalysts to harness natural metabolic cascades for electrochemical applications.^1^ Early work has demonstrated the feasibility of using sequential enzymes, such as invertase and glucose oxidase, to electrochemically monitor sucrose concentration.^2^ More recently, the nitrogenase enzyme complex has been applied to electrosynthesis and the methanol oxidation pathway has been applied to biofuel cells. ^3–5^ These advances highlight the broad potential of bioelectrocatalysis in sensing, electrosynthesis, and energy harvesting.

Beyond simple linear pathways described above, more complex metabolic networks such as the Krebs cycle have been explored for bioelectrocatalytic applications. The Krebs cycle is a cyclical pathway that begins when pyruvate is converted to acetyl-CoA, then combines with oxaloacetate to form citrate. Through a series of oxidation reactions, citrate undergoes stepwise transformation to generate reducing equivalents (NADH) while ultimately regenerating oxaloacetate, which completes the cycle and enables continuous processing of new pyruvate molecules (**Figure S1**).^6,7^ By integrating Krebs cycle enzymes onto a bioanode, pyruvate can be oxidized, electrons can be extracted, and current can be generated.^8^ Importantly, the inclusion of multiple dehydrogenases allows for multiple electron extractions per molecule, enabling deeper oxidation. Previously, a biofuel cell capable of pyruvate oxidation was developed that showed increased power output with each additional dehydrogenase, peaking when the full complement of Krebs cycle enzymes was present.^8–10^

Despite their potential, integrating enzymatic cascades onto bioelectrodes remains challenging. For example, comprehensive modeling of the methanol oxidation pathway revealed multiple engineering parameters that impact a bioelectrochemical system’s efficiency. These parameters not only include individual enzyme kinetics, but the transport of intermediate between enzyme active sites and efficient cofactor regeneration.^11^ Therefore, addressing these challenges requires an integrated approach combining enzyme engineering with material design to translate natural metabolic pathways into efficient bioelectrocatalytic devices.

One significant challenge for enzymatic biofuel cells is the continuous supply of fuel and cofactors to sustain long-term catalysis. The use of NAD^+^-dependent enzymes presents an economic hurdle, as stoichiometric amounts of this relatively expensive cofactor must be added with substrates, and the poor electrochemical performance of NADH on carbon paper electrodes limits its effectiveness as a direct mediator.^12^ To make these devices energetically and economically viable, redox polymers have been integrated onto electrodes to regenerate NAD^+^ electrochemically. These polymers are functionalized with redox-active groups covalently bound to their backbone and facilitate electron transfer via self-exchange reactions.^12^

Protein immobilization is also important for electrode reuse and the elimination of purification steps after solution replenishment. Redox polymers based on linear poly(ethylenimine) (LPEI) have been successfully employed to immobilize various proteins. These LPEI backbones can be crosslinked to form hydrogel matrices that retain enzymes near the electrode surface while permitting substrate and cofactor diffusion. Various mediators, including ferrocene, quinones, and TEMPO, have proven effective in redox catalysis.^13–16^ However, for efficient and selective oxidation of the enzymatic cofactor NADH, diaphorase can be additionally entrapped within the redox polymer, catalyzing the reduction of redox mediators by oxidizing NADH back to NAD^+^. Electrochemical regeneration using diaphorase with redox polymers offers a sustainable approach that minimizes side reactions while maintaining high regioselectivity under mild conditions compatible with most proteins.^17^

Another challenge with bioelectrochemical systems is ensuring efficient intermediate transfer between sequential enzymes while minimizing diffusion losses. In some cascades, nature has addressed this through substrate channeling, an evolutionary strategy that enhances metabolic efficiency by directing intermediates between active sites without releasing them into the bulk solution.^18,19^ Substrate channeling can be achieved by organizing active sites in close proximity and employing structural mechanisms such as intramolecular tunnels, electrostatic guidance, and covalent tethering.^20–22^ Enzymatic cascades that employ substrate channeling have increased catalytic flux, prevent intermediate loss, and drive thermodynamically unfavorable reactions forward.^23–25^

Substrate channeling can occur through intramolecular tunnels, electrostatic guidance, spatial clustering of active sites, and covalent tethering of intermediates.^19^ Two well-characterized examples illustrating distinct strategies are the bifunctional enzymes tryptophan synthase and dihydrofolate reductase-thymidylate synthase (DHFR-TS). Tryptophan synthase catalyzes a two-step reaction of indole 3-glycerol-phosphate to tryptophan, with crystallographic data revealing a 2.5 nm-long hydrophobic tunnel connecting its active sites.^20^ Kinetic studies comparing open and blocked tunnel variants show that in the open variant, the intermediate remains sequestered in the tunnel, enhancing flux and enabling efficient channeling.^26^ In contrast, DHFR-TS lacks a tunnel and instead channels by electrostatic guidance. The two active sites, approximately ∼40 Å apart, are connected by a positively charged surface that is thought to guide the negatively charged intermediate between them.^27^ Experimentally, both the tryptophan synthases and DHFR-TS exhibit significant kinetic advantages over monofunctional enzymes, supporting the role of electrostatic guidance in channeling.^28^

Given the advantages of substrate channeling for enhancing flux and yields, researchers have explored strategies to engineer artificial channeling into cascades, including biofuel cells such as enzymes in methanol oxidation.^29^ These approaches range from DNA and protein-templated assembly to bring enzymes in close proximity, to encapsulation strategies to confine enzymes within protein shells or microcompartments, to direct genetically encoded fusions of sequential enzymes with computationally designed linkers that electrostatically guide intermediate transfer^30–35^ However, despite these diverse strategies, constructing a functional, efficiently channeled enzyme system remains a significant challenge. A major limitation in engineering substrate channeling is the difficulty of detecting and quantifying channeling efficiency. Current methods are largely indirect, rely on complex assays, and suffer from low throughput, making it difficult to rapidly iterate designs and optimize key parameters.^19,28,36^

To support growing efforts in bioelectrochemical energy harvesting and the engineering of substrate channeling in catalytic cascades, we developed a high-throughput fluorescent biosensor assay to screen a library of fusions constructs incorporating linker variants between malate dehydrogenase and fumarase to promote efficient substrate electrostatic channeling. From this screen, a promising fusion enzyme was identified and further characterized using traditional channeling assays, confirming its ability to facilitate intermediate transfer. This substrate channeling fusion enzyme was then incorporated onto electrodes and produced higher current densities than non-channeling fusions. These results highlight the potential of this screening technique for bioelectrocatalytic applications. Additionally, preliminary experiments revealed that immobilization of native tetrameric fumarase led to activity loss, suggesting that its quaternary structure may be destabilized upon attachment to electrode surfaces. To address this issue, a novel single-chain form of fumarase was designed using computational protein design tools. Substrate channeling capabilities were preserved when this *de novo* designed enzyme was fused to malate dehydrogenase using the linker identified via the screening assay, demonstrating that the fusion strategy is applicable even when the fumarase domain has a different overall structure.

## RESULTS

### Evaluating the Function of Neutral Enzyme Fusion Construct V0

*E. coli* malate dehydrogenase (MDH) and fumarase were selected as candidates for engineering a substrate channeling pair due to the highly unfavorable equilibrium constant (K_eq_) of the MDH reaction that limits oxaloacetate yield under physiological conditions (**Figure 1A**). Substrate channeling can help overcome this bottleneck by increasing the local concentration of oxaloacetate and enabling its immediate conversion by fumarase. Notably, there is no natural substrate channeling known between these two sequential steps in the Krebs cycle. Both enzymes are well-characterized and can be readily expressed and purified. Of the three fumarase isoforms in *E. coli*, class I fumarases (FumA and FumB) were not considered due to containing iron-sulfur clusters that are oxygen and temperature sensitive. Instead, the class II fumarase FumC was chosen for engineering.^37–39^

**Figure 1.**
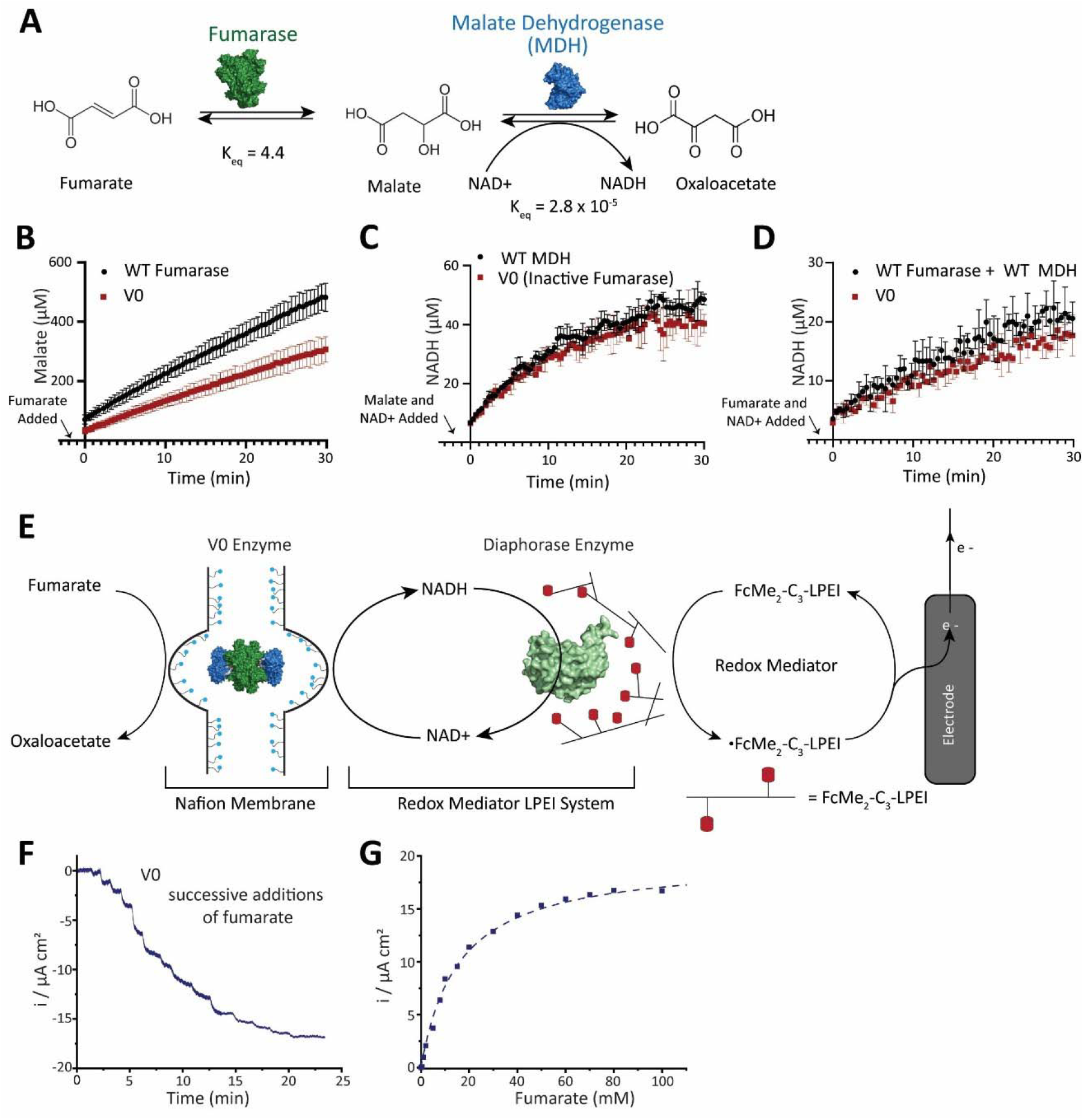
(**A**) Enzymatic cascade of fumarate to malate to oxaloacetate by the enzyme fumarase and malate dehydrogenase (MDH) (**B-D**) V0 in vitro enzymatic activity relative to WT (**B**) V0 Fumarase activity (**C**) MDH activity (**E**) and the ability to turnover fumarate to NADH and oxaloacetate. (**E**) Nafion Electrode Schematic (**F**) V0 Amperometric i-t curve (**G**) V0 relative current density

To promote substrate channeling between MDH and fumarase, a genetically encoded fusion approach was chosen to enable rapid optimization and screening. Based on prior studies showing that a designed cationic linker can facilitate electrostatic channeling of a negatively charged intermediate, we wanted to apply this strategy to an MDH-fumarase fusion.^29^ Electrostatic channeling relies on minimizing the distance between active sites and creating a favorable electrostatic environment for oppositely charged intermediates to travel between them.^40,41^ Given the proximity of the MDH C-terminus and fumarase N-terminus to their respective active sites, the fusion was designed to link these two regions.

A control fusion named V0 was created by linking MDH to fumarase using a flexible, neutral glycine-serine linker (G_4_SG_4_S)^42^ (**Figure S2, S3**) to evaluate if the fusion disrupted protein folding or catalytic activity. Since fumarase functions as a tetramer with active sites at subunit interfaces, fusions to MDH could potentially interfere with proper protein assembly.^37^ *In vitro* enzymatic assays showed that while the MDH portion of the V0 fusion retained full catalytic efficiency, the fumarase component exhibited a 13-fold decrease in catalytic efficiency relative to wild-type (**Figure 1B,C, Table 1)**. This activity loss suggests that fumarase is a more sensitive component than MDH in fusion development. Although MDH is a dimer, its active sites are not located at the subunit interface, unlike fumarase.^43,44^ Therefore, the differing fusion sensitivity may reflect the importance of an intact quaternary structure for fumarase activity, whereas MDH tolerates structural perturbation more readily. Despite the decrease in fumarase activity of V0, because MDH activity remains the rate-limiting step in the cascade, the overall catalytic turnover from fumarate and NAD^+^ to oxaloacetate and NADH was largely unaffected (**Figure 1D, Table 1**).

**Table 1.** Kinetic parameters for wild-type fumarase, wild-type MDH, and the H23 fusion were determined for each enzyme component. The fumarase component catalyzed the conversion of fumarate to malate, the MDH component catalyzed the conversion of malate and NAD^+^ to oxaloacetate and NADH, and the combined components catalyzed the conversion of fumarate and NAD to oxaloacetate and NADH. p-values comparing the V0 and H23 to wild-types enzymes were also determined, in addition to the fold change of the V0 and H23 compared to wild-type enzymes

|  | K<sub>m</sub><br>(μM) | k<sub>cat</sub><br>(s<sup>-1</sup>) | k<sub>cat</sub>/K<sub>m</sub><br>(M<sup>-1</sup> s<sup>-1</sup>) | p-value for k<sub>cat</sub>/K<sub>m</sub><br>comparison to Wild-type Enzyme | k<sub>cat</sub>/K<sub>m</sub> Fold Change from<br>Wild-type Enzymes |  |
| --- | --- | --- | --- | --- | --- | --- |
|  |  |  |  |  | Fold-Change | Direction |
| Fumarase Component – fumarate to malate turnover |  |  |  |  |  |  |
| WT Fumarase | 443 ± 101 | 1180 ± 71 | 2.66 x 10<sup>6</sup> ± 0.63 x 10<sup>6</sup> | N/A | N/A |  |
| V0 Fusion | 759 ± 51 | 153 ± 13 | 2.02 x 10<sup>5</sup> ± 0.22 x 10<sup>5</sup> | 0.018 | 13.2 ± 3.4 | Negative |
| H23 Fusion | 934 ± 66 | 23 ± 4 | 2.5 x 10<sup>4</sup> ± 0.5 x 10<sup>4</sup> | 0.014 | 106 ± 33 | Negative |
| MDH Component – malate and NAD<sup>+</sup> to oxaloacetate and NADH turnover |  |  |  |  |  |  |
| WT MDH | 1.30 mM ± 0.15 | 20.6 ± 2.4 | 1.59 x 10<sup>4</sup> ± 0.26 x 10<sup>4</sup> | N/A | N/A |  |
| V0 Fusion | 1.21 mM ± 0.07 | 16.5 ± 0.8 | 1.36 x 10<sup>4</sup> ± 0.08 x 10<sup>4</sup> | 0.45 | 1.69 ± 0.2 | Negative |
| H23 Fusion | 1.22 mM ± 0.06 | 16.5 ± 1.1 | 1.35 x 10<sup>4</sup> ± 0.1 x 10<sup>4</sup> | 0.44 | 1.17 ± 0.2 | Negative |
| Combined Components – fumarate and NAD<sup>+</sup> to oxaloacetate and NADH turnover |  |  |  |  |  |  |
| WT Unlinked<br>Enzymes | 1.08 ± 0.17 | 4.5 ± 0.6 | 4160 ± 850 | N/A | N/A |  |
| V0 Fusion | 0.97 ± 0.10 | 3.7 ± 0.3 | 3800 ± 500 | 0.73 | 0.1 ± 0.3 | Negative |
| H23 Fusion | 0.93 ± 0.12 | 4.1 ± 0.4 | 4400 ± 700 | 0.83 | 1.06 ± 0.27 | Positive |

Since fumarase and MDH are oligomeric enzymes and are natively found in the *E. coli* expression *strain*, we considered that the observed V0 activity may result from co-purification with endogenous wild-type enzymes forming mixed oligomers. To confirm that the activity originated solely from the engineered fusion protein, V0 was purified from Δ*fumC* or Δ*mdh E. coli* strains. No difference in fumarase or MDH activity was observed for V0 fusions purified from these knockout strains, indicating that the functional V0 complex consists of the fusion protein, without appreciable contribution of native enzymes (**Figure S4**). Given the minimal impact of the fusion on NADH production and the lack of wild-type contamination, V0 serves as a functional control for a fusion lacking substrate channeling and was used as the baseline throughout this study.

### Electrochemical Integration of V0 MDH-Fumarase fusion

A redox polymer system is required to regenerate the NAD^+^ cofactor for the enzyme fusion to perform multi-turnover reactions on the electrode. Two commonly used redox polymers, naphthoquinone-modified linear poly(ethylenimine) (NQ-LPEI) and dimethylferrocene-modified LPEI (FcMe_2_-C_3_-LPEI),^15,16^ were tested for their ability to mediate NADH oxidation by diaphorase, initially without the V0 fusion. Chronoamperometry and cyclic voltammetry showed that NQ-LPEI had poor catalytic performance, while FcMe_2_-LPEI supported significantly higher oxidative current densities. Ferrocene-based electrodes also showed good stability over repeated cycling (**Figure S5A-B**). However, co-entrapment of the V0 fusion protein with diaphorase in either redox polymer severely impaired catalytic activity (**Figure S5C-D**). Although oxidative current could still be triggered by direct NADH addition, highlighting that the diaphorase/polymer layer remained functional, only minimal current was observed with fumarate addition. This data suggests that the polymer matrix itself may compromise V0 activity.

To address this incompatibility, a bilayer electrode architecture was tested that physically separated Vo and the redox polymer. V0 was entrapped in a Nafion matrix modified with tetrabutylammonium bromide (TBAB) and deposited as a second layer over the FcMe_2_-C_3_-LPEI/diaphorase film (**Figure 1E**). Nafion is a micellar polymer that provides hydrophobic pockets suitable for protein encapsulation, while TBAB neutralizes its sulfonic groups and enhances protein compatibility.^17^ This bilayer strategy successfully preserved V0 activity, as clear increases in oxidative current were observed upon fumarate addition, confirming enzymatic turnover and effective NADH regeneration (**Figure 1F-G)**. Therefore, the protective TBAB-Nafion layer offers a means to preserve enzyme function while immobilizing the enzyme and maintaining electrochemical connectivity. These results demonstrate that while redox polymers like FcMe_2_-LPEI support cofactor recycling, their direct interaction with sensitive proteins such as V0 can be detrimental. Since the Nafion-FcMe_2_-LPEI system enabled functional fumarate oxidation on the electrode, it was used in subsequent electrochemical experiments.

### Substrate Channeling Design and Screening

To promote channeling between fumarase and MDH, fusion proteins incorporating a cation-dense linker (KKRQKKKRK) were considered.^29^ This rigid linker is designed to form an α-helix and facilitate electrostatic channeling. However, we hypothesized that the linker alone would be insufficient. Prior studies demonstrate optimal enzymatic proximity is within 10 nm, with channeling efficiency rapidly declining at greater distances.^41^ The active sites of fumarase and MDH are located near their termini but will likely remain in suboptimal distance and orientation if simply linked with the cationic linker. Close active site proximity was achieved previously through enzyme truncation followed by fusion with the cationic linker,^29^ however, a similar truncation with MDH reduced activity (**Figure S6**). As MDH is the rate-limiting enzyme in the cascade and directly impacts maximum electrode current, further activity loss was a concern. Therefore, alternative approaches were required to bring the active sites closer while preserving MDH functionality.

A scaffolding strategy was explored by extending the C-terminus of MDH and the N-terminus of fumarase with flanking scaffolds around the KKRQKKKRK linker. We hypothesized that a properly designed and strategically positioned linker scaffold would maintain correct enzyme folding and improve spatial organization of the cationic linker to promote electrostatic channeling between active sites (**Figure 2A-B**). Given the challenges of precise computational design, including the complexity of predicting dynamic protein interfaces and optimal linker conformations for a tetrameric assembly, a random library-directed evolution approach was taken.

**Figure 2.**
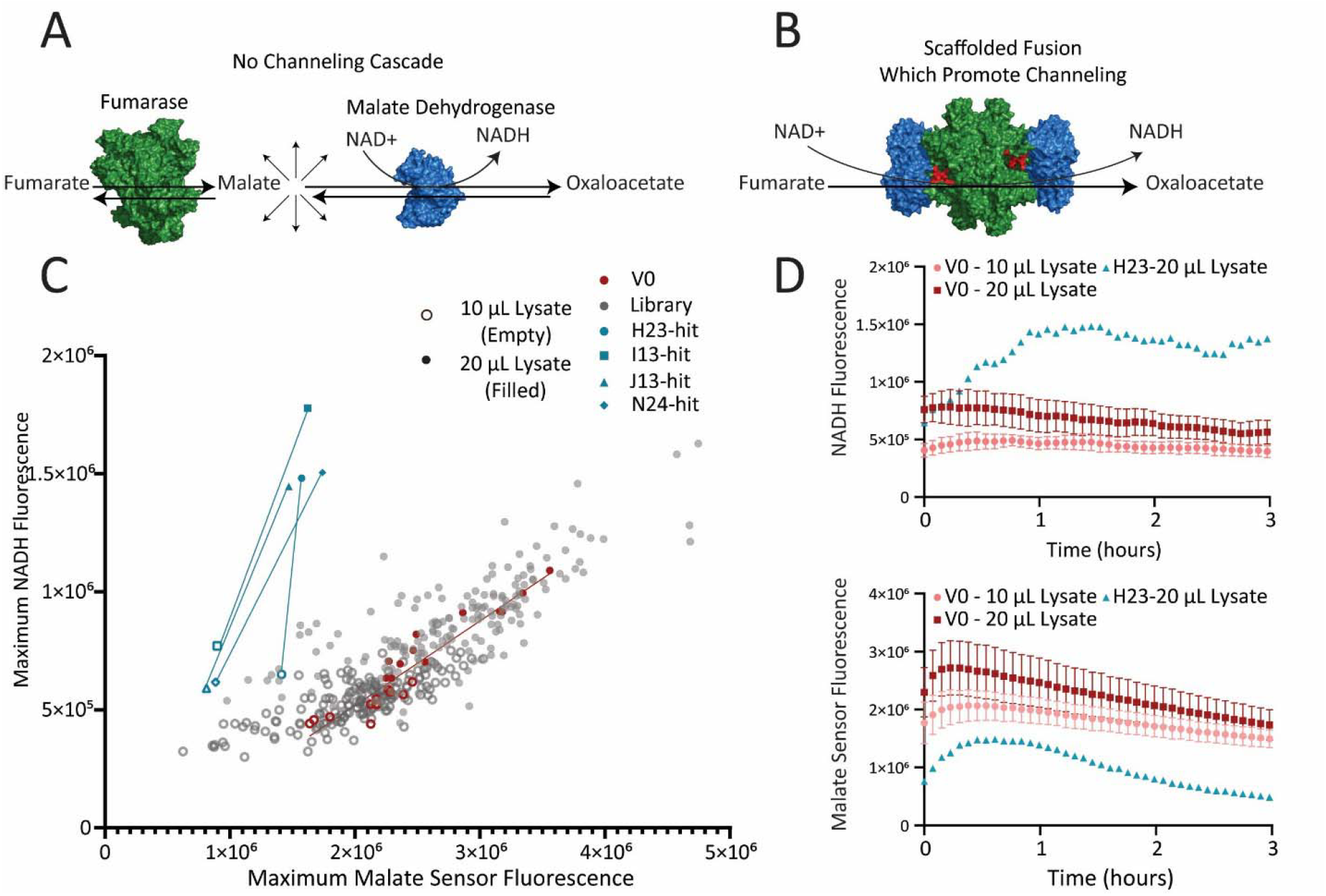
(**A**) Conversion of fumarate to oxaloacetate without channeling (**B**) Conversion of fumarate to oxaloacetate with channeling by properly oriented fusions between fumarase and MDH (**C**) High-throughput screen for substrate channeling - maximal malate sensor signal plotted vs the maximal NADH signal for each colony in the lysate screen (**D**) Malate and NADH curves plotted over time for V0 compared to H23, one of the identified hits.

A fusion construct library utilizing the KKRQKKKRK linker flanked by two randomized eight-residue scaffolds was generated and screened in *E. coli* cell lysate. Scaffolds were constructed using the NDT degenerate codon, which avoids stop codons while providing 12 amino acids representing all major biophysical properties.^45^ To screen for substrate channeling, the real-time conversion of fumarate and NAD^+^ to the intermediate malate and the product NADH was monitored in cell lysates using a recently developed malate fluorescent biosensor^46^ and the intrinsic fluorescence of NADH, respectively (**Figure 2C, Figure S7**). We found that analyzing at least two lysate dilution volumes permitted the determination of an activity “slope” value, which was more informative and less affected by sample-to-sample variations in cell growth, protein expression, and lysis efficiency. While the majority of library members exhibited similar activity slope values to the V0 control, four constructs were identified that exhibited a higher activity slope, corresponding to a higher ratio of NADH to malate production (**Figure 2D**). To ensure reproducibility, the screen was independently repeated on a separate day (**Figure S8**), yielding consistent results across both runs.

Of the four identified hits (**Figure S9**), only the H23 construct was successfully purified for further characterization (**Figure S10**). The other three hits were not successfully purified and instead produced highly fragmented proteins when observed on an SDS gel. Interestingly, H23 exhibited an 8-fold decrease in fumarase catalytic efficiency compared to V0, and a 106-fold decrease relative to wild-type fumarase, primarily due to the fumarase k_cat_ value being lowered to match the MDH k_cat_ value, which is unchanged between H23 and V0 (**Table 1**). In retrospect, screening for less leakage of the intermediate malate also could bias toward fusions that have impaired fumarase activity, as that, in addition to substrate channeling, would reduce malate accumulation in the bulk solution. It is even more notable then, that the apparent catalytic efficiency of the combined activity for H23 appears to be comparable or slightly better than V0 and the unlinked enzymes. Although the improvement is not statistically significant, it highlights that MDH remains the rate-limiting step in this two-enzyme cascade, and that improving substrate channeling to the rate-limiting enzyme can be advantageous even at the expense of upstream steps, as long as they do not become rate-limiting.

### *In Vitro* Biochemical Characterization of Substrate Channeling

Enzymatic cascades can fall into three categories for substrate channeling: no channeling, perfect channeling, or leaky channeling. Perfect channeling is characterized by enhanced product formation with no detectable intermediate. In contrast, leaky channeling occurs when substrate transfer is evident from established *in vitro* protocols but intermediates still accumulate in bulk solution. Initial *in vitro* assays of H23 rule out perfect channeling, as malate is measurable in solution (**Figure S11**). Interestingly, the *in vitro* result with purified H23 is more modest than that observed when screening in lysate. This discrepancy may stem from the *in vitro* experiments containing solely the MDH-fumarase enzyme fusion, whereas the *E. coli* lysate contains all of the Krebs cycle enzymes. These downstream enzymes may amplify a modest leaky channeling effect by further processing oxaloacetate, resulting in higher overall NADH production through le Chatelier’s principle.

To more directly determine whether H23 is substrate channeling, two established *in vitro* biochemical assays were employed: cascade resistance to reaction inhibitors and transient time analysis.^28^ In other multienzyme complexes with characterized channeling, the intermediate remains localized at the active sites, which confers resistance to competitive inhibition.^19,28^ Citrate is a known competitive inhibitor of MDH, so its effect on H23, V0, and the two unlinked enzymes was compared (**Figure 3A-C**).^47^ Whereas no difference was observed between V0 and the unlinked enzymes, as expected, citrate inhibition was reduced for the H23 cascade, with an apparent K_m_ for citrate 5-fold greater than V0.

**Figure 3.**
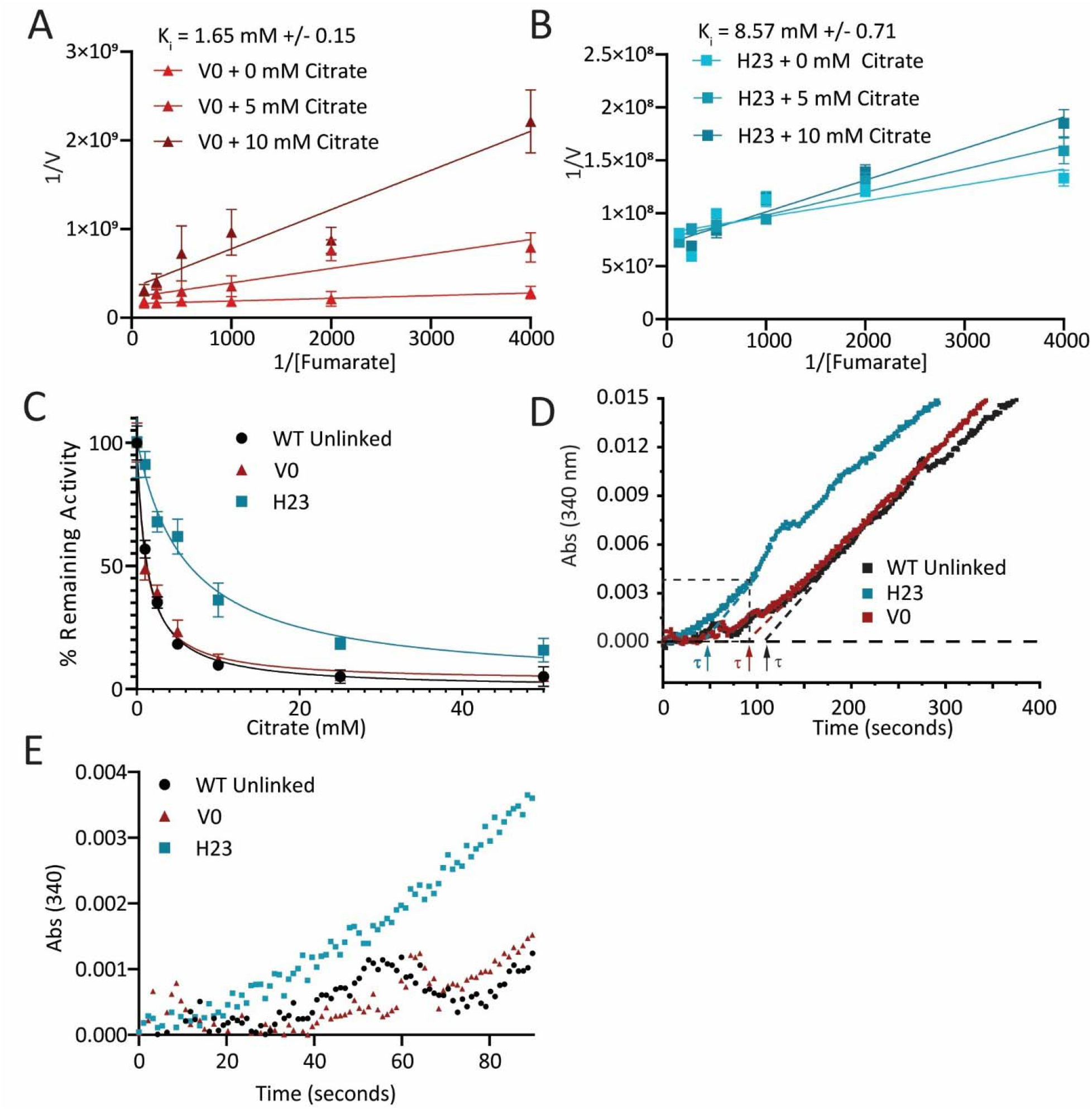
In vitro characterization of substrate channeling with the H23 construct. (**A**) Lineweaver-Burke plot of V0 with varying amounts of citrate (**B**) Lineweaver-Burke plot of H23 with varying amounts of the citrate (**C**) Relative activity of V0, H23 and unlinked WT enzymes with increasing citrate in their ability to turnover fumarate to NADH (**D**) Transient time analysis of H23 vs V0. Dashed box inset shows (**E**) the transient time analysis first 90 seconds

A second method to evaluate substrate channeling is stopped-flow transient time analysis. The lag or transient time (τ) represents the time required to reach steady-state flux of the reaction intermediate. Efficient channeling is expected to reduce τ by accelerating intermediate transfer between active sites.^19,28^ Experiments at varying enzyme concentrations revealed that only low concentrations (<10 nM) produced a measurable delay in NADH production and more pronounced lag times. This concentration-dependent behavior likely reflects increased intermolecular distances at lower enzyme concentrations, minimizing inter-complex malate diffusion and allowing clearer resolution of intra-complex channeling effects.

In time course experiments monitoring NADH production by its UV absorbance, H23 exhibited a significantly shorter τ (43.6 ± 17.8 s) compared to V0 (95.7 ± 26 s) and the two unlinked enzymes (99.5 ± 19.5 s) (**Figure 3D, E**). In contrast, direct injection of malate into the reactions produced no measurable change in τ, with both H23 and V0 showing overlapping activities (**Figure S12**), indicating that the observed lag differences arise specifically from processing of fumarate and not from malate turnover alone. Taken together, these results support that the H23 scaffolded linker promotes substrate channeling and is more efficient at malate transfer between active sites than the V0 flexible neutral linker, which functions no better than the two unlinked enzymes.

### Effect of Substrate Channeling on Product Yields

Beyond traditional lag time and inhibitor studies, substrate channeling should enhance product yield. After allowing the enzymes to equilibrate for 24 hours with fumarate and NAD^+^, H23 demonstrated a 29% increase in NADH production compared to both unlinked enzymes and the V0 control, in addition to an 8% decrease in malate (**Figure 4A-B, Table S1**). To validate that the observed equilibrium differences were due to substrate channeling and not any other effects of the scaffolded linker on MDH activity in the fusion constructs, control experiments were conducted on fumarase-inactive mutants of H23 and V0.^48^ To analyze MDH activity alone, malate was added as the substrate instead of fumarate. No increase in NADH product yield was observed in either fusion construct under these conditions (**Figure S13**), confirming that the enhanced product yield in H23 is due to substrate channeling rather than other beneficial effects from the scaffolded linker.

**Figure 4.**
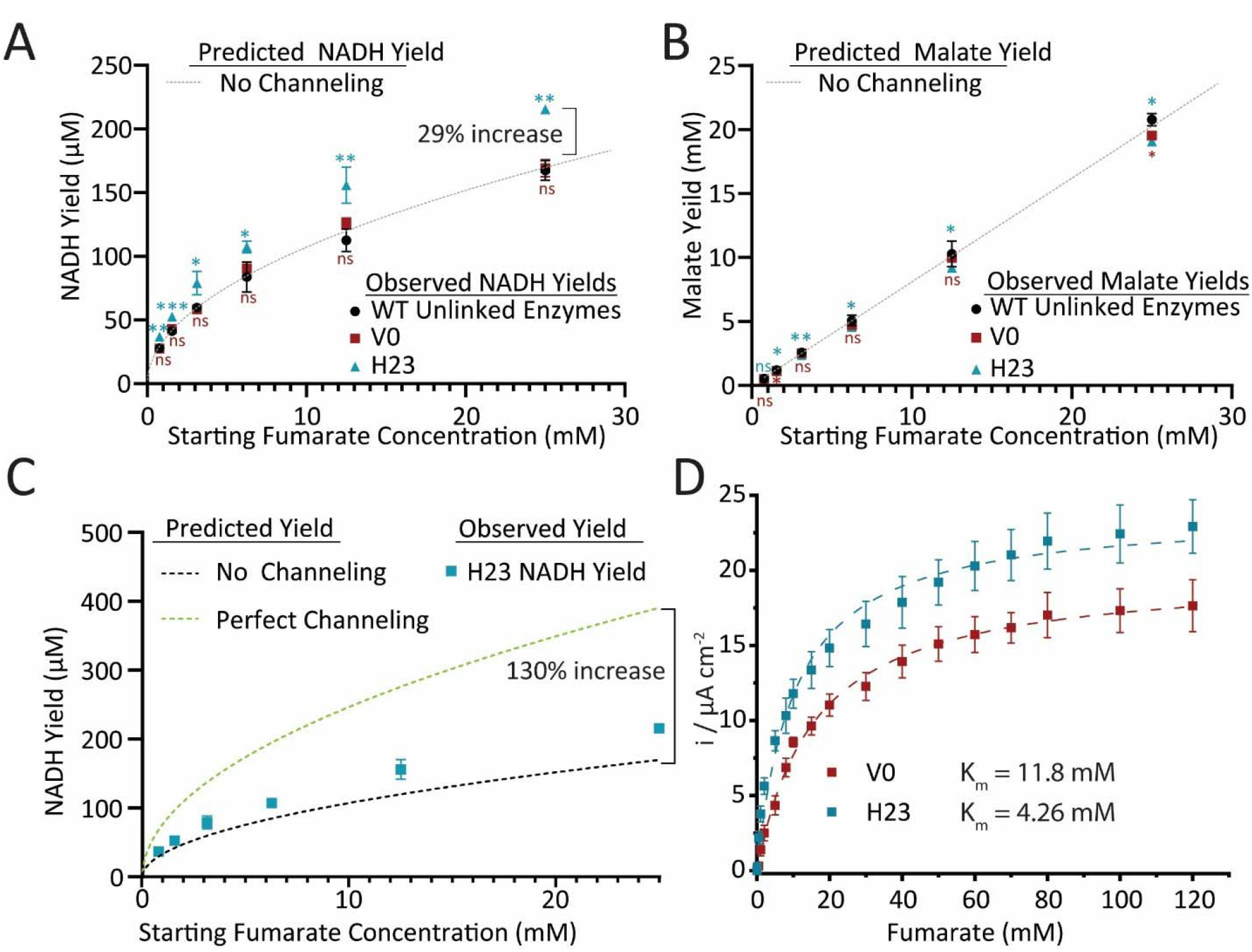
Comparison of equilibrium metabolite yields. Pairwise statistical significance p-values of V0 and H23 yields compared to WT enzyme yields are denoted by stars above or below each set of points (ns = not significant, ^*^ p-value ≤ 0.05, ^**^ p-value ≤ 0.01, ^***^ p-value ≤ 0.001) **(A)** Equilibrium NADH yields of WT enzymes, V0 and H23 compared the predicted NADH from known equilibrium constants **B)** Equilibrium malate yields of WT enzymes, V0 and H23 compared the predicted malate from known equilibrium constants **(C)** H23 NADH yield increase compared to the predicted NADH yield for a no channeling system and a perfect channeling system **(D)** Chronoamperometric comparison of V0 vs H23

Notably for the MDH-fumarase cascade, even under conditions of perfect channeling where no malate leaks into the bulk solution, the maximal theoretical yield increase is only 130%, corresponding to an NADH yield of 388 μM versus the expected equilibrated yield of 169 μM for a starting concentration of 25 mM fumarate. However, with H23, the observed yield was 219 μM. Due to the poor thermodynamics of the cascade, even small changes in NADH can reflect a much larger shift in internal flux (**Figure 4C, see methods and Figure S14**). For H23, the reduction in malate in the equilibrium bulk solution exceeds the corresponding increase in NADH (**Figure S15**), but there is still measurable malate. Thus, H23 appears to be undergoing leaky substrate channeling.

Following confirmation of V0 activity in the bilayer hydrogel system (**Figure 1G**), the same electrode architecture was used to evaluate the H23 fusion. Electrodes containing immobilized H23 exhibited an apparent increase in oxidative current upon sequentially adding fumarate, consistent with NADH generation and oxidation (**Figure 4D**). Compared to V0, H23 generated 29% higher current densities (22.9 ± 1.72 µA cm^-2^ vs. 17.7 ± 1.77 µAcm^-2^), which matches the increased NADH production from fumarate oxidation observed in the solution reactions. Overall, the improved electrochemical response observed for H23 highlights the functional advantage of the H23 protein in the cascade reaction on an electrode.

Interestingly, the electrode-based enzyme kinetics differed markedly from initial solution phase *in vitro* measurements (**Table 1**). Upon immobilization within the electrode polymer, the apparent affinities (K_m_ values) for fumarate decreased for both V0 and H23 to 11.8 mM and 4.26 mM, respectively. While the two enzymes exhibited similar K_m_ values in solution, H23 showed greater affinity for fumarate than V0 when immobilized on the electrode. H23 also displayed a higher current density on the electrode, even though both enzymes had near identical solution-phase turnover rates for fumarate to oxaloacetate, a metric that should correspond to current density.^49^ The reduced substrate affinity upon immobilization may result from structural constraints imposed by the hydrogel matrix. As seen previously with reduced catalytic activity in V0 and H23 fusion constructs, polymer entrapment could impair proper oligomerization and restrict conformational flexibility. The improved performance of H23 relative to V0, of both current density and affinity, may stem from its substrate channeling capability. In solution, diffusion is unrestricted, however, entrapped on the electrode, diffusion of malate likely is limited. In this context, H23’s channeling mechanism may reduce intermediate loss between the fumarase and malate dehydrogenase domains, preserving local substrate concentrations. This, in turn, leads to enhanced catalytic efficiency and supports higher current densities.

### Design and Characterization of a Novel Single-Chain Fumarase

The tetrameric structure of the class II fumarase presents challenges for enzymatic fusion, as previously discussed, due to the active site residues being distributed across multiple subunit interfaces. In this tetramer, each of the four active sites is formed by residues from three different subunits (**Figure 5A**).^37,49^ This spatial arrangement may be perturbed when each monomer is linked to malate dehydrogenase or embedded within a polymeric hydrogel matrix. Structural interference could explain the reduced turnover for fumarase observed in both V0 and H23 variants (**Table 1**), as well as the loss of V0 activity when it was embedded into the LPEI polymer (**Figure S5**).^51,52^

**Figure 5.**
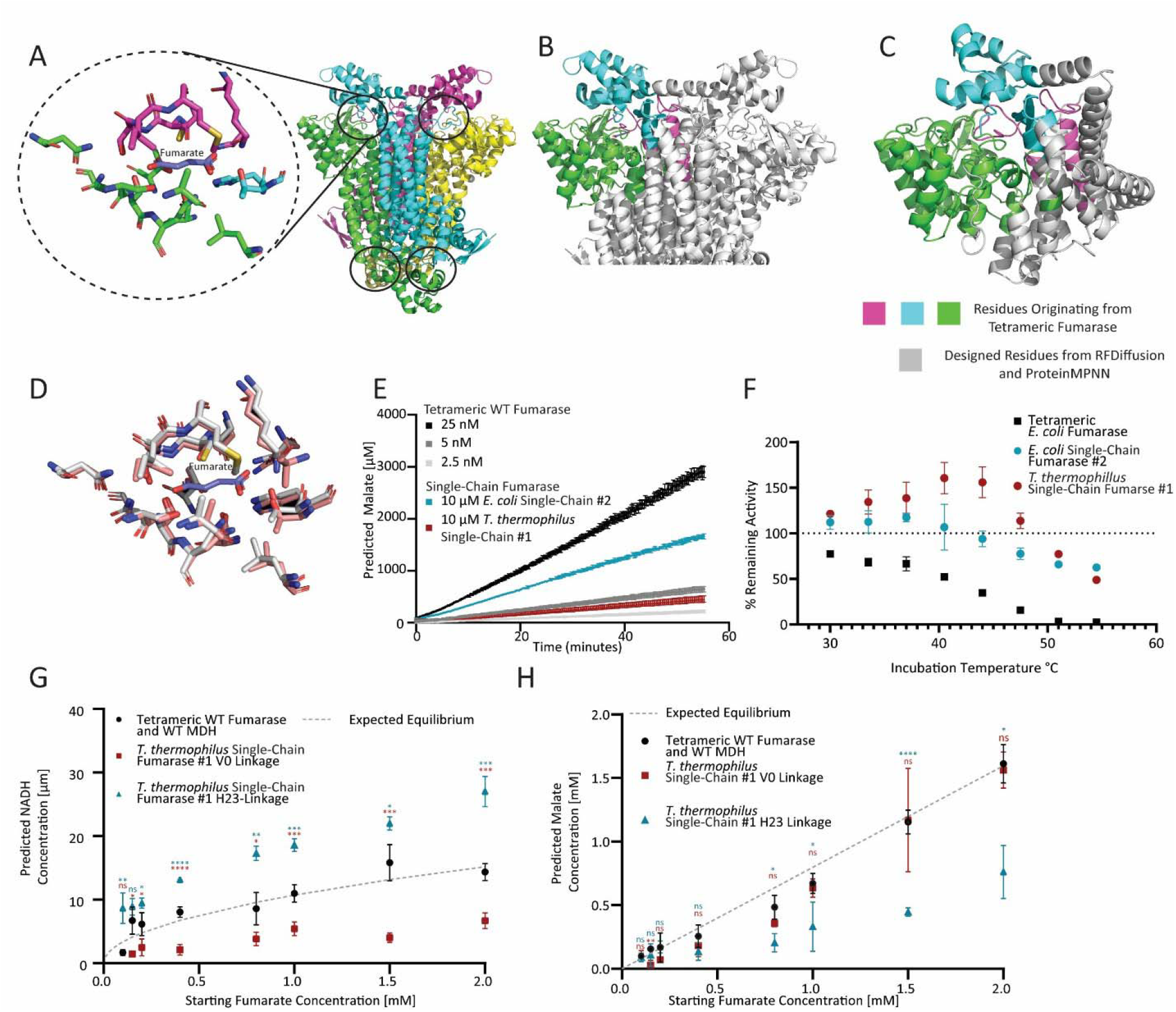
Single-Chain Fumarase design and testing **(A)** Left: Zoomed-in view of a single active site of tetrameric fumarase (PDB: 4APB). Right: Full structure of tetrameric fumarase, with each subunit shown in a different color. The four active sites are circled. **(B)** WT tetrameric fumarase with key residues fragments surrounding the active site used in single-chain fumarase construction colored, discarded residues are colored in grey **(C)** Single-chain Fumarase structure predicted by Alphafold2. Residues originating from WT fragments colored. Residues designed by Rfdiffusion and ProteinMPNN in grey **(D)** Active site of tetrameric fumarase (residues within 10 Å of fumarate) in grey aligned with the predicted active site of Single-chain fumarase by Alphafold2 (colored). Fumarate from the original structure in colored in blue **(E)** Turnover of fumarate to malate by WT tetrameric E. coli fumarase compared to the best functioning single-chain proteins based on E. coli and T. thermophilus fumarase **(F)** Relative activity of WT tetrameric E. coli fumarase compared to both single-chain proteins after heat incubation **(G)** Equilibrium NADH concentrations of WT enzymes, and T. thermophilus single-chain linked to MDH using both the V0 and H23 linker compared the expected NADH from known equilibrium constants. Pairwise statistical significance p-values of single-chain V0 and H23 yields compared to WT enzyme yields are denoted by stars above or below each set of points (ns = not significant, ^*^ p-value ≤ 0.05, ^**^ p-value ≤ 0.01, ^***^ p-value ≤ 0.001 **(H)** Equilibrium malate concentrations of WT enzymes, and T. thermophilus single-chain linked to MDH using both the V0 and H23 linker compared the expected malate from known equilibrium constants

To address the structural instability of tetrameric fumarase, a computational approach was pursued to design a single-chain variant. The active site of class II fumarase is formed by non-contiguous backbone fragments contributed by three subunits, necessitating a strategy to spatially reconnect these elements within a single polypeptide chain while preserving catalytic function. The x-ray crystal structure of the *Mycobacterium tuberculosis* class II fumarase tetramer (PDB 4APB) provided a complete view of the active site architecture and served as the initial reference. From this structure, six discrete backbone segments, spanning multiple chains and positioned within 20–30 Å of the active site, were posited as essential for the formation of the active site (**Figure 5B**). Preserving this spatial arrangement maintains key features such as proton relay networks, which are critical for enzymatic activity.^53^

X-ray crystal structures of *E. coli* (PDB 1YFE) and *T. thermophilus* (PDB 1VDK) fumarases, both class II enzymes, also were selected as design templates due to their relevance for our engineered fusion and enhanced thermal stability, respectively. However, these structures contain only the single subunit form and not the full tetrameric assemblies. To approximate their native quaternary structures, single-chain fumarase designs were aligned to the 4APB tetramer using Pymol, enabling identification of homologous backbone segments that constituted one active site for these fumarases. These catalytic fragments for *E. coli* and *T. thermopilus* fumarases were retained in the modeled structure and all other residues were removed. RFdiffusion was then used to generate *de novo* protein backbones that linked the catalytic segments into a contiguous chain with the aim of maintaining the 3D arrangement of the fumarase active site (**Figure 5C–D**).^50^ Final amino acid sequences were designed using ProteinMPNN to form the target fold, which should preserve the native catalytic geometry within a minimal, single-chain scaffold.^52^

Using this combination of RFdiffusion and ProteinMPNN, a total of 200 candidate sequences predicted to fold as single-chain fumarases were generated. Structural prediction for each designed sequence was generated using AlphaFold2 then aligned to the respective starting tetrameric models to identify the most promising designs based on the lowest RMSD values. From the 200 candidate sequences, three single-chain candidates based on *E. coli* fumarase and three more based on *T. thermophilus* fumarase were selected (**Table S2, Figures S16 and S17)**. These six designs were subsequently expressed and purified for experimental validation. The relative activity of the designed single-chain fumarases was determined by measuring fumarate to malate conversion using the malate biosensor. All six designed constructs exhibited measurable catalytic activity (**Figure S18**); however, even the most active variants showed drastically reduced catalytic efficiency. The best-performing *E. coli-*based variant was 1,886-fold less active than wild-type fumarase, while the top *T. thermophilus* variant was 6,333-fold less active at room temperature (**Figure 5E, Table 2**). However, we found that the designed single-chain fumarase enzymes exhibit increased thermal stability relative to the WT tetramer. Both single-chain enzymes retain full activity up to 44 °C, whereas the tetrameric WT enzyme showed 77% of enzymatic activity at 30 °C, the first temperature tested, and was almost completely inactive by 5 °C (**Figure 5F, Figure S19**). Intriguingly, whereas the single-chain fumarase based on the *E. coli* enzyme retained its activity to higher than room temperature, the *T*. thermophilus-based enzyme showed maximal activity (156 +/- 14.0 % relative to rt) at 40 °C.

**Table 2.** Kinetic parameters of two single-chain Fumarases, E. coli single-chain #2 and T. thermophilus single-chain #1. Additionally, kinetic parameters of the T. thermophilus single-chain #1 linked to MDH using both the V0 and H23 linkage

|  | Fumarase |  |  | MDH |  |  |
| --- | --- | --- | --- | --- | --- | --- |
| | $K_m$ ($\mu\text{M}$) | $k_{\text{cat}}$ ($\text{s}^{-1}$) | $k_{\text{cat}}/K_m$ ($\text{M}^{-1} \text{s}^{-1}$) | $K_m$ (mM) | $k_{\text{cat}}$ ($\text{s}^{-1}$) | $k_{\text{cat}}/K_m$ ($\text{M}^{-1} \text{s}^{-1}$) |
| Single-chain *E. coli* Fumarase #2 | $758 \pm 192$ | $1.07 \pm 0.06$ | $1410 \pm 366$ | N/A | N/A | N/A |
| Single-chain *T. thermophilus* Fumarase #1 | $809 \pm 59$ | $0.34 \pm 0.03$ | $420 \pm 50$ | N/A | N/A | N/A |
| Single-chain *T. thermophilus* Fumarase #1 V0 MDH Linkage | $969 \pm 61$ | $0.33 \pm 0.03$ | $340 \pm 40$ | $1.05 \text{ mM} \pm 0.39$ | $21.6 \pm 1.4$ | $2.06 \times 10^4 \pm 7.8 \times 10^3$ |
| Single-chain *T. thermophilus* Fumarase #1 H23 MDH Linkage | $972 \pm 62$ | $0.33 \pm 0.03$ | $340 \pm 40$ | $1.15 \text{ mM} \pm 0.37$ | $19.8 \pm 1.3$ | $1.72 \times 10^4 \pm 5.7 \times 10^3$ |

The catalytic efficiency (*k*_cat_/K_m_) dropped 13- and 106-fold when tetrameric fumarase was fused to MDH with the V0 and H23 linkers, respectively (**Table 1**). We had hypothesized that the drastically lower fumarase *k*_cat_ values were due to both fusions perturbing fumarase tetramerization. In support of this idea, fusing MDH to the *T. thermophilus*-based single-chain fumarase using either the V0 or H23 linkages did not change fumarase *k*_cat_ values and exhibited only a 1.2-fold reduction in *k*_cat_/K_m_ with the same linkers (**Table 2**). To rule out trace contamination with WT fumarase and confirm that the observed activity arises from the designed single-chain fumarase, mutations that disrupt fumarase catalytic activity^48^ were introduced into the *T. thermophilus* single-chain MDH-fumarase fusion. Both active and mutated fusion enzymes were purified and tested for fumarase activity, which showed that malate production required the active single-chain fumarase, so was not due to background activity (**Figure S20**). Unfortunately, these same experiments could not be performed with the more active *E. coli*-based single-chain fumarase because the fusion construct was found to be insoluble so could not be purified.

The scaffolded cationic linker H23 facilitated substrate channeling from the wild-type tetrameric fumarase to MDH. Promisingly, increased product yield also was observed when the *T. thermophilus* single-chain MDH-fumarase fusion bearing the H23 linker was incubated with fumarate and NAD^+^ for 48 h to ensure reaction equilibration. The H23-linked single-chain fusion showed a 63% increase in NADH product yield (**Figure 5G, Table S1**) compared to the unlinked enzymes (with WT tetrameric fumarase) that produced the expected amount of NADH at equilibrium. In contrast, the V0-linked single-chain fusion showed lower NADH product yield. Furthermore, the intermediate malate was reduced by 55% for H23-linked single-chain fusion relative to unlinked enzymes (**Figure 5H**). These shifts in product and intermediate distribution are substantially greater than those observed for the H23 construct with WT tetrameric fumarase, which showed a 29% increase in NADH production and an 8% decrease in malate intermediate.

Taken together, these results demonstrate that computational protein design can be applied to improve the stability of an enzyme that forms a sensitive oligomer, converting it to a single polypeptide chain. While the designed enzymes were orders of magnitude less active than the wild-type tetramer, reflecting the limitations of modeling the active site geometry based on ground state structures rather than the transition state, their activities were less affected by the fusion to the linker and MDH. In addition, the H23 linker, which was originally screened for substrate channeling between the WT tetrameric fumarase and MDH, could be directly applied and exhibited more efficient substrate channeling from the designed single-chain fumarase to MDH.

## CONCLUSION

In this work, we demonstrate the first engineered example of substrate channeling between fumarase and malate dehydrogenase via a genetic fusion approach. Using a dual-fluorescence screening assay that simultaneously reports on intermediate and product levels, we rapidly evaluated libraries of fusion constructs in crude cell lysates. To our knowledge, this represents the first screening campaign specifically designed to evaluate channeling using separate fluorescence signals for both the intermediate and product. This dual screening approach enabled the rapid evaluation of genetically encoded enzyme variant libraries and improved upon traditional assays that require multiple indirect kinetic experiments. More broadly, this approach could be applied to DNA and protein-templated engineering strategies during the design phase, allowing researchers to rapidly test variations in scaffold length, topology, or attachment sites to optimize intermediate transfer efficiency. These advances offer a streamlined path for optimizing multi-enzyme pathways, with implications for metabolic engineering, cell-free bioprocessing and bioelectrochemical systems where efficient intermediate transfer is critical.

We developed analysis steps in the dual-fluorescence screen to overcome variability in cell growth and protein expression, extracting trends even in complex lysates. The screen also revealed important design principles: selecting for higher product-to-intermediate ratios unexpectedly enriched for variants with reduced fumarase activity, underscoring that balanced catalytic rates are as critical as spatial proximity for efficient channeling to be observed. Although the design and limited availability of analogous biosensors for other metabolite pairs remain as potential bottlenecks, this work establishes a general framework for high-throughput channeling evaluation that can be extended to diverse enzymatic pathways, which can accelerate the development of enzymatic-derived biofuel cells.

Beyond establishing a novel substrate channeling assay, our screen identified H23 as a fusion construct that promotes intermediate transfer between fumarase and MDH. H23 exhibited channeling in biochemical assays, yielded higher NADH production compared to controls, and produced 29% greater current densities when immobilized on electrodes. These results confirm that screening for altered product-to-intermediate ratios can successfully yield functional channeling variants and that such designs translate into measurable electrochemical advantages. Together, this validates biosensor-guided screening as a practical route toward catalytic biomaterials with enhanced performance.

The enhanced metabolic flux and higher current output observed for H23 pave the way for its integration in artificial cascades that mimic the Krebs cycle metabolon in biodevices, which are characterized to have improved catalytic efficiencies. H23 is of particular interest for the fabrication of pyruvate, ethanol, and lactate bioanodes with increased degree of substrate oxidation and number of electrons generated.^12^ The present work has undertaken the challenge of creating a functional enzymatic design where the channeling efficiency of H23 is coupled with the continuous generation of NAD+ cofactor for sustainable energy conversion. The presented bioelectrode design allows for the future incorporation of other NAD+ dependent dehydrogenases and enzymes in the modified Nafion membrane, coupling the resulting multi-enzymatic cascade with the redox polymer-diaphorase-based NAD+ regeneration system on the electrode.

This work also demonstrates the first successful computational redesign of tetrameric fumarase into functional single-chain variants. By preserving the essential active site architecture, we generated enzymes that retained measurable catalytic activity while showing enhanced thermal stability and tolerance to genetic fusion. Importantly, the single-chain fumarases with the H23 substrate-channeling linker yielded 63% higher NADH production compared to unlinked controls. These results establish a strategy for converting sensitive oligomeric enzymes into more robust single-chain forms, broadening the applications of the resulting single-chain enzymes in synthetic biology and bioelectrochemical applications

The designed enzymes were orders of magnitude less active than wild-type fumarase, underscoring persistent challenges in computational enzyme design.^54^ As the method used relied on ground-state structures and did not adequately capture transition-state geometries or active-site dynamics, this likely contributed to the reduced catalytic efficiency. Additionally, low purification yields of designed constructs limited classical substrate channeling assays and electrode integration. Nevertheless, the improved stability and fusion tolerance of the single-chain variants make them promising starting points for directed evolution aimed at recovering activity while preserving their advantageous properties.

An important takeaway from this study is the role of kinetic balance between linked enzymes. When the first enzyme in a cascade is significantly faster than the second, intermediates are more likely to diffuse away, reducing the observed channeling. In the tetrameric H23 construct, channeling coincided with reduced fumarase turnover, better matching the downstream MDH activity (**Table 1**). Similarly, with even slower turnover, the single-chain enzymes yielded higher product accumulation, potentially due to a better aligned reaction rate with MDH. These findings underscore that effective channeling designs must optimize spatial arrangement and catalytic rates between pathway enzymes.

For future research, this work highlights several promising directions for advancing substrate channeling in cell-free bioprocessing and bioelectrochemical applications. Fluorescence-based detection of intermediates and products offers a platform for high-throughput screening of channeling configurations and could be extended to other metabolic cascades where suitable biosensors are available or can be developed. Within the Krebs cycle, future optimization should focus on enhancing the kinetics of both MDH and the single-chain fumarase, as MDH remains the rate-limiting step and limits overall pathway flux and current density. The enhanced thermal stability and fusion tolerance of the single-chain fumarases make them attractive starting points for directed evolution, which could recover catalytic activity while preserving these advantageous properties. Continued exploration of linker design and enzyme configurations will also be critical, as both spatial proximity and kinetic balance have now been shown to influence cascade efficiency. Together, these directions highlight how computational design, biosensor-guided screening, and directed evolution may converge to enable programmable control of multi-enzyme assemblies.

### Experimental Section

#### Safety Statement

No unexpected or unusually high safety hazards were encountered.

#### Cloning Methods

##### General Transformation Protocol

A 1 μL aliquot of plasmid at a concentration of at least 20 ng/μL was added to a 25 μL aliquot of chemically competent frozen XL1-Blue *E. coli* (UC Berkeley Macrolab, Berkeley, CA, USA). This was done in 1.5 mL Eppendorf tubes. The cell plasmid mixture was incubated on ice for 30 min. The cells then underwent a heat shock procedure: tubes containing the cells and plasmid were added to a heat block set to 42 °C for 45 secs then immediately returned to ice for 2 min. 300 μL of sterile LB broth was then added, and cells were recovered for 45 min in an incubator set at 37 °C with shaking at 250 rpm. After recovery, 50 μL of transformed cells were plated onto an LB agar plate containing 50 μg/mL of carbenicillin. The plates were allowed to incubate at 37 °C overnight or for 16-18 hrs.

##### General Plasmid Construction

A two-piece Gibson assembly methodology was used to generate plasmids encoding protein variants, with all constructions performed within the pBAD plasmid backbone. The base plasmid was obtained from the Citron biosensor plasmid (Addgene #134303), which featured an N-terminal His tag, T7 gene 10 leader sequence, and Xpress tag. The His-tag component was used for subsequent protein purification processes. The complete leader sequence preceding the protein insertion site comprised the peptide ‘MGGSHHHHHHGMASMTGGQQMGRDLYDDDDKDPSSR’, which includes the affinity tag and other molecular components.

Polymerase chain reactions were performed using Phusion DNA polymerase (UC Berkeley Macrolab) in a total reaction volume of 50 μL. The reaction mixtures consisted of 29.5 μL sterile water, 10 μL of 5x Phusion HF buffer, 2.5 μL each of 10 μM forward and reverse primers, 1 μL of template DNA at approximately 10 ng/μL, 5 μL of 2.5 mM dNTPs, and 0.5 μL of Phusion polymerase. The thermocycling protocol began with an initial denaturation at 98 °C for 30 secs, followed by 34 cycles of amplification. Each cycle included denaturation at 98 °C for 30 secs, primer annealing for 30 secs at a temperature specifically calculated for each primer pair using the NEB Temperature Calculator, and extension at 72 °C for 150 secs. The reactions were completed with a final extension step at 72 °C for 5 min.

DNA amplification was confirmed through gel electrophoresis using a 2% agarose gel, with 10 μL of the PCR product loaded for visualization. The remaining 40 μL of DNA underwent enzymatic digestion by incubating with 1 μL of Dpn1 restriction enzyme (New England Biolabs) at 37 °C for 1 hr. Following digestion, the DNA was purified utilizing Qiagen PCR cleanup columns following manufacturer’s instructions to remove enzymes, primers, and other reaction components. The final DNA sample was quantified and assessed for purity using a Thermo Scientific NanoDrop 8000.

For each Gibson assembly, two PCR reactions were typically conducted, with each primer pair designed to contain overlapping regions between the two PCR products to facilitate ligation (Tables S3 and S4). Plasmid mutations were introduced by encoding specific changes within the primer overhangs. Proteins such as fumarase and malate dehydrogenase (MDH) were amplified using genomic *Escherichia coli* DNA as the initial template. For single-chain fumarase construction, the sequence was first processed through the Integrated DNA Technologies (IDT) codon optimization program, with the optimized sequences then gblock fragments ordered from IDT. A primer pair was then designed to amplify the pBAD plasmid backbone, incorporating overhangs matching the first and last 20 base pairs of the IDT-synthesized product.

Gibson assembly reactions were prepared by combining 5 μL of pre-combined PCR products or for single-chain fumarase plasmids, the PCR backbone and resuspended IDT-ordered DNA with 5 μL of NEB 2× Gibson assembly master mix. Multiple DNA ratios were tested, including 1:2, 2:1, and 1:1, with total DNA concentrations ranging from 0.02 to 0.05 pmol. The reaction mixture underwent assembly by incubation at 50 °C for 1 hr in a thermal cycler.

The 10 μL of PCR/Gibson mix was transformed into XL1-Blue *Escherichia coli* cells using the previously described transformation protocol. Following plating on LB agar containing 50 μg/mL of carbenicillin, 2-3 bacterial colonies were isolated and cultured overnight in 3 mL of sterile LB broth supplemented with 50 μg/mL of carbenicillin. Plasmid DNA was subsequently extracted from the overnight cultures using Qiagen miniprep columns and verified through whole plasmid sequencing (Plasmidsaurus).

##### V0 Plasmid Construction

The V0 plasmid was constructed through a two-step Gibson assembly process involving malate dehydrogenase (MDH) and fumarase C genes from XL1-Blue *E. coli*. The initial phase involved amplifying MDH from the *E. coli* genome and cloning it into the pBAD plasmid using Gibson assembly, creating a foundational plasmid capable of MDH expression. A subsequent round of Gibson assembly integrated the fumarase C gene.

Two distinct PCR amplifications were used in this process. The first reaction used the newly created MDH-containing plasmid as a template, with primers designed to bind to specific regions of the plasmid. The forward primer was bound immediately after the MDH gene, while the reverse primer bound to the MDH gene’s 3’ end, effectively linearizing the entire plasmid. Simultaneously, a second PCR reaction amplified the fumarase C gene directly from *E. coli* genomic DNA.

The primer was designed to incorporate the glycine-serine linker (GGGGSGGGGS). The reverse primer of the fumarase C amplification contained overhangs corresponding to the region directly following the MDH gene, while both the fumarase forward primer and the plasmid backbone reverse primer included the glycine-serine linker sequence. These designed overhangs served as overlap regions during the Gibson assembly, allowing for the integration of the fumarase gene into the MDH-containing plasmid.

##### Inactivating Fumarase Mutation

Inactive fumarase mutants for V0, H23, and single-chain fumarase were made by introducing several alanine mutations in the active site: S318A, S319A, K324A, and N326A. Previous research, combined with the x-ray crystal structure (PDB code 4APB) assessment of fumarate bound in the active site demonstrated that these residues are important for catalysis and substrate binding^37,50^. Mutagenesis was performed using a two-piece Gibson assembly, as described above, with PCR amplification of the original plasmids as templates (Tables S3 and S4).

##### Cationic Linker Library Construction

To promote substrate channeling between fumarase and malate dehydrogenase (MDH), fusion proteins were constructed that incorporated a rigid cation-dense linker (KKRQKKKRK) flanked by randomized scaffolds to position the cationic linker between active sites. The V0 construct was the starting PCR template, as it contained the MDH and fumarase genes already linked, with the C terminus of MDH linked to the N terminus of fumarase by the glycine serine linker. For library construction, two separate PCR reactions were performed on the V0 template plasmid. The first PCR used a forward primer that annealed to the plasmid backbone and a reverse primer binding to the 3’ end of the MDH gene. This reverse primer contained an overhang of 24 random nucleotides (8 NDT codons) followed by the KKRQKKKRK sequence. The second PCR used a forward primer binding to the 5’ region of the fumarase gene with an overhang consisting of the KKRQKKKRK sequence followed by 24 random nucleotides (8 NDT codons). The reverse primer for this reaction annealed to the plasmid backbone near the forward primer of the first PCR pair, creating an overlap region for subsequent Gibson assembly.

The KKRQKKKRK sequence in both primer overhangs served as a common overlap region for the Gibson assembly, in addition to the backbone region used in both primers. The resulting PCR products were combined by Gibson assembly as described above. The Gibson mixture was transformed into chemically competent *E. coli* XL1-Blue cells and plated on LB agar containing 50 μg/mL of carbenicillin. Each resulting colony contained a different variant of the scaffold-linker-scaffold configuration between the two enzymes. While this approach generated a library with a theoretical size of 10^17^ unique fusion constructs, only ∼2000-4000 colonies were observed across 4 plates, and only ∼800 colonies were screened.

#### Fluorescence-Based Lysate Screen

To screen the cationic linker library, approximately 800 colonies were individually picked and grown overnight for ∼16 hrs in 300 μL of LB + carbenicillin at 37 °C with shaking at 250 rpm in sterile 96 DeepWell plates (ThermoFischer Scientific). In addition to the scaffolded fusion protein plasmid, this screen was also performed on a control plasmid V0, containing MDH linked to fumarase with a flexible glycine-serine linker. Following overnight growth, 30 μL of each overnight culture was subcultured into fresh 270 μL LB + carbenicillin + 0.1 % arabinose to induce the expression of the fusion proteins from the pBAD plasmid. Additionally, 200 μL of sterile 50 % glycerol solution was added to the original overnight cultures, which were then stored at −80°C. The fresh subcultures were grown overnight at 37 °C with shaking at 250 rpm in sterile 96 DeepWell plates.

The following morning, the overnight cultures were pelleted by centrifugation for 10 min at 5000 RCF. After discarding the LB supernatant, cell pellets were resuspended in 300 μL lysis buffer (100 mM potassium phosphate buffer, pH 7.5, 0.4 % Triton X-100, 1 mg/mL lysozyme). Resuspended cells were incubated in lysis buffer for 1 hr at 4 °C to ensure complete lysis.

For substrate channeling activity screening, variable quantities of cell lysate (10 μL or 20 μL) were transferred to 384-well black-bottom Greiner plates. Variable lysate volumes were used to compensate for differences in cell growth and protein expression between samples. To provide internal reference points for interpreting the screen, both a low-yield (10 μL) and a high-yield (20 μL) lysate sample of the baseline construct (V0) were included on each plate. These controls helped establish a range of expected signal behavior against which screened variants could be compared. To each well containing lysate, 45 μL of reaction buffer (100 mM potassium phosphate pH 7.5, 25 mM fumarate, 50 mM NAD^+^, and 1 μM Malon sensor) was added. Plates were analyzed on a SpectraMax i3x plate reader (Molecular Devices) for approximately 4 hrs, continuously monitoring two fluorescence signals: NADH (340/460 nm ex/em) and Malon (500/540 nm). The maximal RFU recorded for Malon and NADH over the 4 hrs of monitoring were plotted against each other (Figure 2C). These two signals allowed for the visualization of the production of the intermediate malate (via Malon)^46^ and the final product NADH. To compensate for variations in cell growth and expression levels, the ratio of NADH product formed to the intermediate malate was analyzed.

Constructs with consistently high NADH/Malon production ratios between the two replicate screens were identified as potential substrate channeling hits. Non-induced cell cultures frozen at −80 °C with glycerol were retrieved, and 10 μL were added to 3 mL fresh LB + carbenicillin and grown overnight. Overnight cultures were pelleted at 5000 RCF for 10 min. Plasmids were then extracted from the cell pellet using the Qiagen Qiaprep columns for plasmid purification. Whole plasmid sequencing (Plasmidsaurus) was performed to identify the scaffolded sequences.

#### General Protein Purification Protocol

Plasmids containing the desired protein sequence in the pBAD backbone were transformed into Xl1-Blue cells and plated on LB agar plates supplemented with 50 μg/mL carbenicillin. After overnight growth, a single colony containing the sequence-confirmed construct was picked with sterile toothpicks and grown overnight in a sterile 12 mL culture tube containing 3 mL of LB media supplemented with 50 μg/mL carbenicillin in a 37 °C incubator, shaking at 250 rpm. After overnight growth in 3⍰mL of LB media, the overnight culture was added to a 3 L baffled flask with 1 L of 2x YT media and 50 μg/mL carbenicillin. This large culture was grown until the OD reached ∼0.5-0.8 (typically 3-6 h), then a sterile 10% arabinose water solution was added to a final concentration of 0.1% to induce protein expression. The induced culture was incubated in a shaker at 30 °C, 250 rpm, for 16 hrs.

After overnight induction, the cells were collected by centrifugation at 5000 RCF for 30 min at 4 °C. The spent media was discarded, and the cell pellet was resuspended in 50 mL of lysis buffer (50 mM Tris-HCl, 300 mM NaCl, 5 % glycerol, 10 mM imidazole, pH 8.0) with the addition of 10 mg lysozyme (Millipore Sigma) and 150 μL of 200 mM PMSF dissolved in ethanol (Sigma Aldrich). The cell lysate was transferred to a 50 mL falcon tube and sonicated for 5 min, with 2 s pulses on and 2 s off with a 70% amplitude using a sonicator (Fisherbrand). During sonication, the falcon tube containing the cells was immersed in an ice-bath to maintain a low temperature. The sonicated cells were centrifuged again at 5000 RCF for 40 min at 4 °C to obtain clarified lysate.

The clarified lysate was decanted into a new 50 mL falcon tube and incubated with 2 mL of HisPur Nickel NTA resin (Thermo Scientific) for 1 h with rotation at 4 °C. The lysate and resin mixture were added to an Econo-Pac 20 mL chromatography column (Bio-Rad) to collect the resin in the column and remove the flow-through. The nickel resin was washed with ∼60 mL of wash buffer (50 mM Tris-HCl, 500 mM NaCl, 25 mM imidazole, 25% glycerol pH 8.0) followed by elution of bound protein with 8 mL of elution buffer (50 mM Tris-HCl, 500 mM NaCl, 500 mM KCl 250 mM imidazole, 25% glycerol pH 8.0).

The eluted protein was concentrated and the buffer was exchanged with protein storage buffer (50 mM Tris-HCl, 100 mM NaCl pH 7.5 with 10% glycerol) using Amicon Ultra Centrifugal Filter columns with a 30 kDa cutoff with repeated centrifugation at 4000 RCF at 4 °C. The concentration of the final protein was calculated using a Bradford assay (Thermo Scientific Pierce BCA Protein Assay Kit) following the manufacturer’s instructions.

Protein purity was analyzed on a Bio-Rad TGX FastCast Acrylamide gel. Briefly, 20 µL of protein sample was mixed 1:1 with loading buffer (250 mM Tris-HCl pH 6.8, 8% w/v SDS, 0.02% w/v bromophenol blue, 200 mM DTT) and heat denatured at 95 °C for 5 min. Denatured protein mixtures (20 µL) were loaded into cast wells of polyacrylamide gels, and 2 µL of prestained protein ladder (NEB Color Prestained Protein Standard, Broad Range, 10–250 kDa) was included in the first lane. Gels were run in Tris–glycine– SDS running buffer (25 mM Tris base, 192 mM glycine, 0.1% SDS) at a constant 160 V at 4 °C for ∼60 min, or until the dye front reached approximately three-quarters of the gel length. Following electrophoresis, gels were rinsed in water, immersed in staining buffer (Coomassie Brilliant Blue R-250 0.1% w/v in 50% methanol, 10% acetic acid) for 30–45 min, and destained in buffer (50% methanol, 10% acetic acid) until protein bands were clearly visible. After purification, protein samples were stored in 10-20 μL aliquots at −80 °C. Prior to use, protein aliquots were thawed at room temperature, kept on ice, and discarded after 1 freeze-thaw cycle.

For single-chain fumarase purification, slight adjustments were made to the purification protocol. The 1 L of 2x YT media was supplemented to final concentrations of 200 mM arginine, 300 mM sorbitol, and 0.3% (v/v) glycerol. Lysis buffer, wash buffer, and elution buffer also were supplemented to a final concentration of 300 mM arginine, 300 mM sorbitol and 25% (v/v) glycerol added in addition to the reagents previously mentioned.

#### Electrochemical Experiments

##### Preparation of Tetrabutylammonium bromide/Nafion membrane

For the preparation of the modified Nafion, a 3-fold molar excess of tetrabutylammonium bromide (TBAB; Sigma Aldrich) was added to 2 mL of a 5% (w/v) Nafion suspension (Ion Power, Inc.). The mixture was vortexed at 1100 rpm for 15 min and then poured into a plastic weighing tray (3 inch × 3 inch). The solvent was allowed to evaporate slowly, leaving behind a transparent, light-yellow film at the bottom of the tray. The resulting film was soaked in 20 mL of 18 MΩcm deionized water for 24 hrs to extract any excess of unreacted alkyl ammonium bromide and HBr, followed by multiple water rinses. After the complete evaporation of residual water under ambient conditions, a transparent, plastic-like membrane formed. This membrane was resuspended in 2 mL of a 1:1 ethanol/water mixture containing 3 ceramic beads and stored at 4 °C.

##### Preparation of redox polymer/DH and TBAB-Nafion/fusion proteins electrodes

NQ-LPEI and Fc-LPEI redox polymers were synthesized as previously reported. Stock solutions of the redox polymers, diaphorase (DH; AG Scientific, Inc., 1 KU), and the crosslinker ethylene glycol diglycidyl ether (EGDGE; Polysciences, Inc.) were prepared in Milli-Q water. Carbon paper electrodes (Fuel Cell Earth, Inc.) were fabricated according to the method of Gerulskis et al.^55^ In brief, commercial carbon paper (Fuel Cell Earth, Inc.) was cut into 3 cm X 0.5 cm individual stripes using a craft-cutting machine. Subsequently, the stripes were immersed for a few seconds into melted wax and then cooled down, leaving an uncovered electroactive surface area of 0.25 cm^2^ on one end of each stripe. For the preparation of the electrodes with diaphorase entrapped in the NQ-LPEI redox polymer, 35 μL of 10 mg/mL NQ-LPEI, 15 μL of 3 mg/mL DH, and 2.5 μL of 10% (v/v) EDGDE were gently mixed. A total of 15 μL of the mixture was deposited onto the 0.25 cm^2^ delimited area of each electrode. The electrodes were then left to cure for 6 hrs at 25 °C to promote the crosslinking process before carrying out electrochemical measurements. The electrodes containing DH entrapped in the Fc-LPEI redox polymer were prepared similarly by mixing 40 μL of 10mg/mL FcMe_2_-C_3_-LPEI, 30 μL of 3 mg/mL DH, and 2.5 μL of EDGDE 10% v/v. A 15 μL aliquot was deposited on the electrode surface and cured under the same conditions. To immobilize fusion proteins on the electrodes via entrapment in the TBAB-Nafion film, either 13 μL of V0 or H23 protein (170 μM) was mixed with 6.5 μL of the modified TBAB-Nafion suspension. A final volume of 12 μL of this mixture was deposited on the delimited area of the electrode previously modified with either NQ-LPEI/DH or FcMe_2_-C_3_-LPEI/DH layer. The electrodes were then kept at 4 °C for 2 hrs before measurements.

##### Electrochemical experiments (characterization of polymers with fusion proteins)

All electrochemical experiments were performed using a CH Instruments potentiostat in a standard three-electrode setup with a platinum mesh counter electrode and a saturated calomel electrode (SCE) as the reference. Modified carbon paper electrodes were immersed in potassium phosphate buffer (pH 7.4) for 30 min to allow swelling and polymer equilibration. Electrooxidation of NADH by NQ-LPEI/DH and FcMe_2_-C_3_-LPEI/DH was assessed by cyclic voltammetry and chronoamperometry. A potential of +0.25 V vs SCE was applied in chronoamperometric experiments in stirred buffer solution by recording the current generated upon the addition of increasing concentrations of NADH at room temperature. Chronoamperometric experiments with electrodes containing fusion proteins immobilized in the TBAB-Nafion film were carried out at +0.25 V vs SCE in the stirred phosphate buffer solution containing 20 mM NAD^+^. After establishment of a stable baseline, sequential injections of 1M fumarate disodium salt solution (from 100 nM to 140 mM) were carried out in the electrochemical cell. After each fumarate addition, a new baseline was established before proceeding with the next addition, obtaining current-time staircase plots. Plots of the current density at the steady-state values versus fumarate concentration allowed the determination of Michaelis−Menten parameters for each fusion protein via non-linear regression analysis (Origin software version 2023b).

#### De Novo Design of Single-Chain Fumarase

The available structures of *E. coli* and *T. thermophilus* fumarase (PDB 1YFE and 1VDK, respectively) are monomeric and not bound to fumarate. Therefore, the x-ray crystal structure of *Mycobacterium tuberculosis* fumarase (PDB 4APB) as a tetramer with bound fumarate was used as the starting template. When a single *E. coli* fumarase subunit (1YFE) was aligned with each subunit in the tetrameric *M. tuberculosis* structure, the RMSD was 0.832 +/- 0.024 Å. Similarly, aligning a single *T. thermophilus* fumarase subunit (1VDK) with the *M. tuberculosis* structure yielded an RMSD of 1.143 +/- 0.041 Å. In comparison, the four subunits of the *M. tuberculosis* fumarase align to each other with an average RMSD of 0.090 +/- 0.018 Å. Given the structural similarity between the *E. coli, T. thermophilus*, and *M. tuberculosis* fumarases, four subunits from each were aligned with the tetrameric *M. tuberculosis* structure to approximate the tetrameric forms of these enzymes. These tetrameric approximations of *E. coli* and *T. thermophilus* were then used for single-chain design.

Retrospective modeling using AlphaFold3 generated tetrameric fumarase structures for *T. thermophilus* and *E. coli*. Sequences were submitted to the AlphaFold Server (https://alphafoldserver.com/) with the entity type set to “protein,” the copy number specified as four, and a randomly generated seed. For each input sequence, AlphaFold3 produced five predictive models. The predicted tetrameric models were aligned to the approximate tetrameric models used for downstream protein design. The AlphaFold3 structures showed high alignment with the approximated models, with RMSD values of 0.293 ± 0.04 Å for *E. coli* and 0.3162 ± 0.023 Å for *T. thermophilus*. Although the AlphaFold3 models were not directly used in design, they served as an independent validation of the structural accuracy of the approximated models used in monomeric design efforts

Analysis of the tetrameric fumarase structure (PDB 4APB) identified residues within 20-30 Å of the active site that were essential for tetramer formation. Protein design software RFdiffusion and ProteinMPNN were used to generate structural scaffolds that bridged the homologous fragments in the *E. coli* and *T. thermophilus* approximate models into a single contiguous chain capable of folding into a functional fumarase active site^51,52Alphafold2 was used to predict t^.

The design process involved the selection and linking of specific structural fragments from the approximate tetrameric fumarase structures. Six fragments were identified that contribute to a single active site for the *E. coli* model: residues 1-146 from subunit 1, residues 224-273 from subunit 1, residues 285-332 from subunit 3, residues 352-368 from subunit 1, residues 175-200 from subunit 2, and residues 401-467 from subunit 2. These fragments were connected using glycine linkers of varying lengths (11, 18, 22, 100, and 61 residues, respectively) to create a backbone topology that could potentially accommodate the active site geometry. The software RFdiffusion was used to design these linker regions with conformations that would be theoretically compatible with protein folding^5^. These glycine residues served as spatial placeholders, defining the overall fold without specific sequence determinants. The *T. thermophilus* construct mirrored the *E. coli* design, with minor variation: residues 1-146 from subunit 1, residues 224-273 from subunit 1, residues 285-332 from subunit 3, residues 352-368 from subunit 1, residues 176-201 from subunit 2, and residues 401-466 from subunit 2. Glycine linkers: 11, 20, 22, 100, and 60 residues, respectively.

The software ProteinMPNN was then used to replace the glycine placeholders with optimized amino acid sequences predicted to stabilize the designed fold^4^. This two-step computational approach allowed the creation of a single-chain protein that theoretically contained all the structural elements necessary for fumarase activity within a single polypeptide chain.

Using this computational design protocol, 20 distinct structural topologies for single-chain fumarases were generated using RFdiffusion, 10 based on *E. coli* fumarase and 10 based on *T. thermophilus* fumarase. While each topology had an identical length of residues connecting fumarase fragments, they differed in spatial arrangement of these linkers. For each structural topology, ProteinMPNN was set to generate 10 unique protein sequences, resulting in a library of 200 potential fumarase single-chain sequences. To separately evaluate the structure of these designed sequences, Alphafold2 was used to predict the structure of each designed sequence, which was then aligned with the 4 active site fragments used in the original model design.^56^ Six sequences, three generated from the *E. coli* fumarase and three from the *T. thermophilus* fumarase, were selected for *in vitro* characterization (Table S2). These six sequences demonstrated the most promising structural characteristics based on their Alphafold2 predictions predictive models having the highest structural similarity (lowest RMSD) to the original active site structure.

#### In Vitro Enzymatic Assays

Enzymatic assays were performed in 100 mM potassium phosphate buffer at pH 7.4 and 1 μM of purified Malon biosensor. *In vitro* analyses were conducted on a SpectraMax i3x plate reader (Molecular Devices) with excitation/emission wavelengths of 500/540 nm for Malon to measure malate concentrations and 340/460 nm to measure NADH concentrations. All experimental samples were analyzed in technical triplicate using Greiner black 96-well or 384-well plates.

Sample concentrations of NADH or malate were determined by comparison to an equation generated from a standard curve. Control wells were prepared containing all starting reagents (such as fumarate or NAD+) without enzymes. These control wells enabled calculation of the ΔRFU/RFU for enzyme-containing wells and for wells with known metabolite concentrations. ΔRFU/RFU is defined as the change in fluorescence relative to baseline, where ΔRFU is the difference between the fluorescence of the sample well and the corresponding control well, and RFU refers to the baseline fluorescence of the control. This ratio was used to normalize signal intensity and account for background fluorescence.

The resulting ΔRFU/RFU values were used to generate a standard curve for each metabolite and derive the two regression equations that relate ΔRFU/RFU values to log[metabolite]. Analysis of kinetic data was performed using the GraphPad Prism10 software. For NADH, a nonlinear regression (log[agonist] vs. response) yielded an equation with an R^2^ value of 0.9899 and applicable over the range of 1 µM to 20 mM. For malate, a linear regression was used as previously described,^46^ with an R^2^ of 0.949 and applicable in the range of 100 µM to 3 mM.

##### General Enzyme Activity Screening

Initial fumarase and MDH enzyme activity screening was conducted using 5 nM or 1 nM purified enzyme respectively. For fumarase screening, fumarate was added to wells containing 1 μM Malon sensor, 100 mM potassium phosphate buffer (pH 7.4) and variable concentrations of enzyme. The final well volume was 150 μL, with final concentration of fumarate at 25 mM. The turnover from fumarate to malate was observed as an increase in fluorescence from the Malon sensor. For MDH screening, no malate sensor was used; instead, 50 mM NAD+ was added along with 25 mM malate. The increasing NADH signal was monitored using the 340/460 channel. For the screening of fusion enzymes, 25 mM fumarate was added to a solution containing 50 mM NAD+, and the NADH signal was monitored.

To determine if H23 exhibited perfect or leaky channeling, the above experiment was replicated with purified H23 compared to purified V0 with 10 nM enzyme. Fumarate was added to wells containing 1 μM Malon sensor, and 50 mM NAD+ in 100 mM potassium phosphate buffer (pH 7.4). The well volume was 150 μL, with the final concentration of fumarate at 25 mM. Both Malon and NADH signals were monitored.

##### Single-Chain Fumarase Activity Screen

To determine if the initial single-chain fumarases had activity, fumarate was added to wells containing 1 μM enzyme (*E. coli* single-chain variants) or 5 μM enzyme (*T. thermophilus* single-chain variants), and 1 μM Malon sensor in 100 mM potassium phosphate buffer (pH 7.4). The final well volume was 150 μL, with the final concentration of fumarate at 25 mM. Malon signal was monitored for 60 min to verify the fumarate to malate turnover (**Figure S8)**. High enzyme concentration was used as the turnover was expected to be lower compared to tetrameric enzymes. The most promising candidates from each class (*E. coli* single-chain #2, and *T. thermophilus* single-chain #1) were analyzed in duplicate along with samples of WT (wild-type) tetrameric *E. coli* fumarase at different dilutions to compare their activities (**Figure 5E**).

To verify that the fumarate to malate turnover was due to enzymatic activity and not background signal, alanine mutations were made into the active site of the single-chain fumarase. In tetrameric fumarase, residues S318A, S319A, K324A, and N326A were mutated to alanine to abolish activity without disrupting structure. The same mutations were applied to the *T. thermophilus* single-chain #1 linked to MDH with the V0 linkage. Both the active and inactive forms of this construct were analyzed under identical assay conditions as described above.

#### Enzyme Kinetics Analysis

For kinetics experiments involving fumarase, MDH, and combined apparent kinetics, a Lineweaver-Burk plot was constructed by plotting the inverse of the maximal reaction rate against the inverse of the initial substrate concentration. Linear regression analysis of this plot allowed for the determination of kinetic parameters, with x and y intercepts used to calculate the Michaelis constant (K_m_) and catalytic rate constant (K_cat_), respectively. Analysis of kinetic data was performed using the GraphPad Prism10 software.

##### Fumarase Kinetics

Fumarase enzyme kinetic parameters were quantified using a Thermo Scientific NanoDrop 8000, using fumarate’s ultraviolet absorption at 230 nm. A standard fumarate calibration curve was established across a concentration range of 10 mM to 100 μM to ensure quantitative measurements. Absorbance properties were used as opposed to the fluorescent malate sensor due to its response mechanism, which requires several min of binding for a fluorescence response to be quantitative.

Reactions were prepared with 20 nM (tetrameric fumarase) or 1 μM (single-chain fumarase) of enzyme and fumarate concentrations ranging from 0.5 mM to 5 mM in potassium phosphate buffer. Immediately upon mixing fumarate and enzyme, 2.5 μL of the reaction mixture was placed on the NanoDrop pedestal, with absorption measured at 230 nm to quantify fumarate concentration. Measurements were taken every 12 secs for approximately 2 min, resulting in 12 total time points.

Fumarate concentrations were calculated for each time point, enabling prediction of malate production by subtracting the current fumarate concentration from the initial concentration. Plotting the predicted malate production against time allowed observation of enzymatic reaction progression. The linear phase of malate production was used to determine the maximal reaction rate at each fumarate concentration. Kinetic parameters were derived by plotting the maximal reaction rate against the initial fumarate concentration, and using Lineweaver-Burk plots as previously described.

##### Malate Dehydrogenase (MDH) Kinetics

MDH kinetic analysis used 1 nM of purified enzyme (pure MDH or fusion variants) in 150 μL of potassium phosphate buffer containing 50 mM NAD+. Fusions of MDH to fumarase such as V0 and H23 had inactivating mutations at the catalytic residue in fumarase to prevent the reverse direction conversion of malate to fumarate^50^ (Table 1). Variable malate concentrations were introduced to reaction wells, with NADH production monitored via fluorescence measurements over 30 min. NADH quantities were calculated using the previously established standard curve, with the linear phase of NADH production used to determine the maximal reaction rate at each malate concentration. Kinetic parameters were derived by plotting the maximal reaction rate against the initial malate concentration, employing the analytical approach described in previous experimental procedures.

##### Combined Apparent Kinetics

Reactions assessing fumarate conversion to oxaloacetate and NADH were conducted in 150 μL of potassium phosphate buffer containing 3 nM of enzymes. WT enzyme systems utilized 3 nM each of MDH and fumarase to maintain equimolar ratios. Fusion enzymes like V0 and H23 were used at 3 nM, with their genetic fusion ensuring an equimolar composition. The reaction mixture was supplemented with 50 mM NAD^+^ and variable fumarate concentrations. Following fumarate addition, NADH was quantified using the established standard curve. The linear phase of NADH production was analyzed to determine the maximal reaction rate at each fumarate concentration. Kinetic parameters were derived by plotting the maximal reaction rate against the initial fumarate concentration, employing the analytical approach described in previous experimental procedures.

##### Knockout Strain V0 purification

To confirm that V0 activity arose solely from the fusion construct and not wild-type contamination, enzyme kinetics were performed on V0 purified from both Δ*fumC* and Δ*mdh* strains from the *E. coli* Keio collection.^57^ To make chemically competent cells, each strain was streaked from glycerol stocks onto LB agar plates supplemented with 50⍰μg/mL kanamycin and incubated at 37 °C for 16 hrs. A single colony was inoculated into 3⍰mL of LB + kanamycin (50⍰μg/mL) in a 12⍰mL culture tube and grown overnight at 37 °C with shaking at 250⍰rpm. The culture was then transferred to 100⍰mL of fresh LB + kanamycin in a 1⍰L baffled flask and incubated at 37 °C, 250⍰rpm, until reaching an OD_600_ of ∼0.4. Cells were chilled on ice and harvested by centrifugation at 4500⍰×g for 10 min at 4 °C. Pellets were washed once in 100⍰mL ice-cold 100⍰mM MgCl_2_, pelleted again, and resuspended in 25⍰mL of ice-cold 100⍰mM CaCl_2_. After incubation on ice for 20 min, cells were centrifuged again and resuspended in 25⍰mL of ice-cold 85⍰mM CaCl_2_ with 15% glycerol. A final centrifugation step was followed by resuspension in 2⍰mL of 100⍰mM CaCl_2_. Aliquots of 100⍰μL were flash frozen and stored at −80 °C.

The V0 plasmid encoding the MDH–fumarase fusion was transformed into chemically competent Δ*fumC* and Δ*mdh* strains using standard heat shock transformation. Transformed cells were plated on LB agar supplemented with 50⍰μg/mL kanamycin and incubated at 37 °C to obtain colonies. Protein expression and purification were performed as described previously, with all LB and 2xYT media supplemented with 50⍰μg/mL kanamycin to maintain plasmid and selection pressure for the knockout background.

For MDH activity, 10⍰nM of purified V0 fusion protein (from both wild-type and Δ*mdh* strains) was assayed in 150⍰μL of potassium phosphate buffer containing 50⍰mM NAD^+^ and 50⍰mM malate. NADH production was monitored via fluorescence over 30 min to assess relative catalytic activity. For fumarase activity, 1⍰nM of purified V0 fusion (from wild-type or Δ*fumC* strains) was incubated in 100⍰mM potassium phosphate buffer (pH17.4) containing 25⍰mM fumarate and 1⍰μM Malon sensor, in a final volume of 150⍰μL. Malon fluorescence was monitored to quantify fumarate-to-malate turnover. The use of the Malon sensor enabled sensitive detection of product formation even at low fumarase activity levels.

##### Citrate Inhibition

Two sets of citrate inhibition experiments were performed to characterize enzyme activity. In the first set, the percentage of remaining enzyme activity at various citrate concentrations under the following standard conditions: 0.5 nM enzyme, 500 μM NAD+, 2 mM fumarate, and 100 mM potassium phosphate buffer (pH 7.4) in 150 μL reaction volumes was measured on a plate reader. Citrate concentrations tested ranged from 0 to 50 mM, specifically including 0, 1, 2.5, 5, 10, 25, and 50 mM. Fumarate and NAD^+^ conversion to NADH was monitored using the plate reader following established methods. The enzyme’s turnover rate (per sec) was calculated using the linear portion of the NADH production curve at each citrate concentration. Relative enzyme activity was determined by comparing the turnover rates at each citrate concentration to the control condition with no citrate addition, and the results were plotted to visualize the inhibition pattern.

In the second set of experiments, the apparent inhibition constant (K_i_) of citrate on the enzymatic cascade was determined. The experimental setup maintained consistent parameters of 0.5 nM enzyme, 500 μM NAD^+^, and 100 mM potassium phosphate buffer. Fumarate concentrations were varied across a wide range: 250 μM, 500 μM, 1 mM, 2 mM, 4 mM, and 8 mM. For each fumarate concentration, multiple citrate inhibitor concentrations were tested (0, 1 mM, 2.5 mM, 5 mM, 10 mM), and NADH turnover was calculated using the previously established method. Lineweaver-Burke plots were generated for each citrate inhibitory concentration to determine the apparent K_i_ through standard kinetic analysis protocols.^58^

##### Lag time experiments

Stopped-flow lag time experiments were performed using a BioTek Synergy HTX multimode plate reader. NADH production was monitored by measuring absorbance at 340 nm immediately following the injection of fumarate into a quartz cuvette containing a quiescent solution. Reactions were carried out in 0.1 M potassium phosphate buffer (pH 7.4) at 25 °C, with 5 mM NAD ^+^ and either fusion proteins (V0 or H23) or a 1:1 mixture of free fumarase and malate dehydrogenase proteins. Fusion proteins, were used at a final concentration of 5 nM (corresponding to 5 nM of each enzymatic component in a 1:1 ratio). For the unlinked enzyme mixture, 5 nM fumarase and 5 nM malate dehydrogenase were added separately to maintain an equivalent 1:1 molar ratio of enzymatic activities, resulting in a total protein concentration of 10 nM. Prior to fumarate addition, protein and NAD ^+^ were equilibrated in the buffer for 10 min. Fumarate was then rapidly injected to a final concentration of 50 mM. The experimental lag time (τ) was determined by extrapolating the tangent of the maximum slope from the normalized absorbance versus time plot back to an absorbance value of 0.

##### Thermal stability

Enzymatic activity was determined by measuring fumarate to malate conversion across a range of temperatures between 30 and 55 °C. Enzymes were prepared by diluting tetrameric fumarases to 5 nM and single-chain fumarases to 2 μM in 100 mM potassium phosphate buffer (pH 7.4), and maintained on ice before temperature-specific treatments.

The thermal stability experiment used 50 μL enzyme aliquots incubated in PCR strip tubes in a Biorad C1000 Touch Thermal Cycler. The temperature range included 30, 33.5, 37.5, 40.5, 44, 47.5, 51, and 54.5 °C, with an 8-minute incubation period. A control was established by maintaining a separate enzyme aliquot on ice throughout the experiment. Following temperature treatment, 40 μL of each enzyme sample was combined with 40 μL of reaction buffer, resulting in a final composition of 25 mM fumarate, 1 μM Malon sensor and 2.5 nM tetrameric fumarase, or 1 μM single-chain fumarase in 100 mM potassium phosphate buffer (pH 7.4).

Fumarate to malate conversion was monitored using a plate reader in a 384-well plate, tracking the reaction over 1 hr. While the Malon sensor is too slow to report on real-time malate formation, relative changes could be analyzed. Control wells without enzyme were used to calculate ΔRFU/RFU, enabling the quantification of relative malate production.

The enzyme’s relative turnover rate (per sec) was measured at each incubation temperature. Relative enzyme activity was assessed by comparing the turnover rates of heat-incubated enzymes to the non-incubated control, with results plotted to illustrate the temperature-dependent activity profile.

#### Enzyme Equilibrium Analysis

Two sets of experiments were conducted to investigate how substrate channeling enzymes influence the equilibrium between fumarate and oxaloacetate. The first set focused on tetrameric enzymes, while the second set examined single-chain enzyme behavior, with WT enzymes serving as a control for both experiments.

For tetrameric fumarase experiments, reactions were performed in 96-well black bottom Greiner plates with 150 μL reaction volume containing 100 mM potassium phosphate buffer (pH 7.4), 80 nM tetrameric enzymes, 50 mM NAD^+^, and varying fumarate concentrations (0.78, 1.56, 3.12, 6.25, 12.5, and 25 mM). To minimize evaporation, plates were sealed with parafilm and wrapped in aluminum foil, then stored at 4 °C for 24 hrs to ensure equilibrium was reached.

Final NADH concentrations were determined by measuring NADH fluorescence values after 24 h using a plate reader, with control wells without enzyme used to calculate ΔRFU/RFU. Additionally, 75 μL of the overnight reaction mixture was combined with 75 μL of 100 mM potassium phosphate buffer containing 2 μM Malon sensor in new 96-well plates. The mixture was incubated for 10 min to allow the sensor to equilibrate with malate in solution, and fluorescence readings were then collected. Malate concentrations were calculated from a standard curve generated under identical incubation and measurement conditions, and values were doubled to account for the 1:1 dilution.

Using the reported equilibrium constants for fumarate to malate (4.4) and malate to NADH (2.86 × 10^-5), the endpoint equilibrium was calculated for each initial fumarate concentration.^38,59^ The calculated endpoint concentrations of NADH and malate were then plotted against their expected concentrations. (Calculations are explained in the next section). For single-chain fumarase, to accommodate the lower protein yield and slower conversion rate, several modifications were implemented. Reactions were conducted in 384-well plates with a reduced 80 μL volume and the 3 single-chain proteins (T1, T1-H23, T1-V0) had increased enzyme concentration to 1 μM (compared to 80 nM for tetrameric enzymes). The reaction time was extended to 72 hrs to ensure complete equilibration. Fumarate concentrations for the single-chain enzyme experiments were significantly lower, ranging from 100 μM to 2 mM (100, 150, 200, 400, 800 μM, 1 mM, 1.5 mM, and 2 mM), and starting NAD^+^ concentrations were at 5 mM as opposed to 50 mM. These adjustments were necessitated by the slower reaction kinetics of single-chain fumarases, allowing for more precise equilibrium measurements with limited starting material.

##### Calculating Equilibrium NADH Concentrations

To determine the endpoint equilibrium concentrations in the sequential reaction cascade from fumarate to oxaloacetate, a system of equations based on mass balance and equilibrium constraints was established. The reactions considered were:

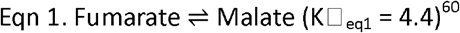

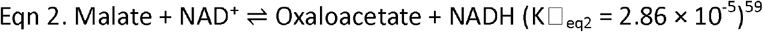

An example calculation is provided for the initial conditions:

Starting Fumarate Concentration: 0.025 M

Starting NAD ^+^ Concentration: 0.05 M

To solve for the equilibrium concentrations, endpoint metabolites were assigned the following variables:

Fumarate = **x**

Malate = **y**

NAD ^+^ = **z**

Oxaloacetate + NADH = **a**

(since they are produced in equivalent amounts **a**/2 = oxaloacetate or NADH)

The following equations were established:

Mass balance:

The total fumarate-derived species must remain 25 mM:

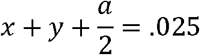

The total NAD species must remain 50 mM:

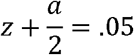

Equilibrium expressions:

For the fumarate ⇌ malate reaction:

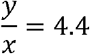

For the malate + NAD ^+^ ⇌ oxaloacetate + NADH reaction:

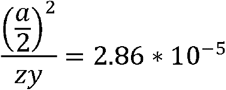

The system of equations was solved to determine the endpoint values. Solving for the variable **a**, and dividing by 2, provides the expected NADH concentration, while solving for **y** gives the expected malate concentration.

To generate an endpoint curve using different initial fumarate concentrations, only the first equation needs to be adjusted:

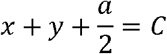

where **C** represents the starting fumarate concentration. By substituting different values for **C**, endpoint concentrations of NADH (**a/2**) and malate (**y**) can be calculated for a range of initial conditions.

##### Theoretically Perfect Channeling Equilibrium Calculations

To estimate the theoretical yield increase under perfect channeling conditions, equilibrium calculations were performed. Using the Gibbs free energy equation,

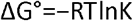

The standard free energy change for the fumarate-to-malate conversion was calculated as ΔG°=−3.67 kJ/mol using an experimentally determined equilibrium constant (K_eq_=4.4).^38^ Similarly, for the reaction

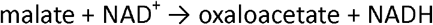

the free energy change was determined to be ΔG°=+25.92 kJ/mol based on an experimentally determined K_eq_=2.86×10^−5^. ^59^ Under perfect channeling conditions, where fumarase is fully coupled to malate dehydrogenase (MDH) with no malate leakage into the bulk solution, the overall reaction simplifies to:

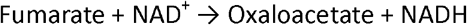

In this scenario, the Gibbs free energy change for the overall reaction is:

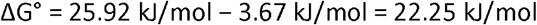

Recalculating the equilibrium constant using this ΔG° yields a theoretical Keq=1.25×10−4. This new equilibrium constant was then used to calculate the expected NADH concentration at an initial fumarate concentration of 25 mM, and the results were compared to the expected NADH levels from the uncoupled reaction.

For the theoretically perfect channeling the equilibrium equations were greatly simplified. The variables were defined as:

Fumarate = x

NAD ^+^ = y

Oxaloacetate + NADH = z

And the following equations were established:

Fumarate-derived species

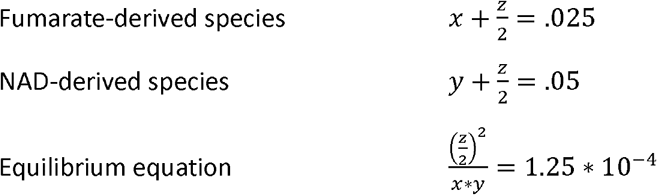

The system of equations was solved simultaneously to determine the endpoint values. Solving for the variable **z**, and dividing by 2, provides the expected NADH concentration.

## Supporting Information

Additional figures are provided in Supporting Information.

## Author Contributions

N.J.R, M.B., and F.A conducted the research, E.B provided synthetic material, N.J.R, M.B, S.D.M., and M.C.H. designed the experiments, N.J.R, M.B., and M.C.H. wrote the manuscript, and S.D.M. and M.C.H. supervised the project. All authors reviewed and edited the manuscript.

## Acknowledgements

This work was supported by the Office of Naval Research (grant number N00014-21-1-2188). The support and resources from the Center for High Performance Computing at the University of Utah are gratefully acknowledged.

