## Supplemental Materials for "Fluorescent Biosensor-Guided Engineering of Enzyme Cascades for Electrochemical Applications"

**TABLE OF CONTENTS**

**Supporting Figures 1-20 and Tables**	pages 3-26

Supplemental Figure 1	page 3

Supplemental Figure 2	page 4

Supplemental Figure 3	page 5

Supplemental Figure 4	page 6

Supplemental Figure 5	page 7

Supplemental Figure 6	page 8

Supplemental Figure 7	page 9

Supplemental Figure 8	page 10

Supplemental Figure 9	page 11

Supplemental Figure 10	page 12

Supplemental Figure 11	page 13

Supplemental Figure 12	page 14

Supplemental Figure 13	page 15

Supplemental Figure 14	page 16

Supplemental Figure 15	page 17

Supplemental Figure 16	page 18

Supplemental Figure 17	page 19

Supplemental Figure 18	page 20

Supplemental Figure 19	page 21

Supplemental Figure 20	page 22

Supplemental Table 1	page 23

Supplemental Table 2	page 23

Supplemental Table 3	page 24

Supplemental Table 4	page 26

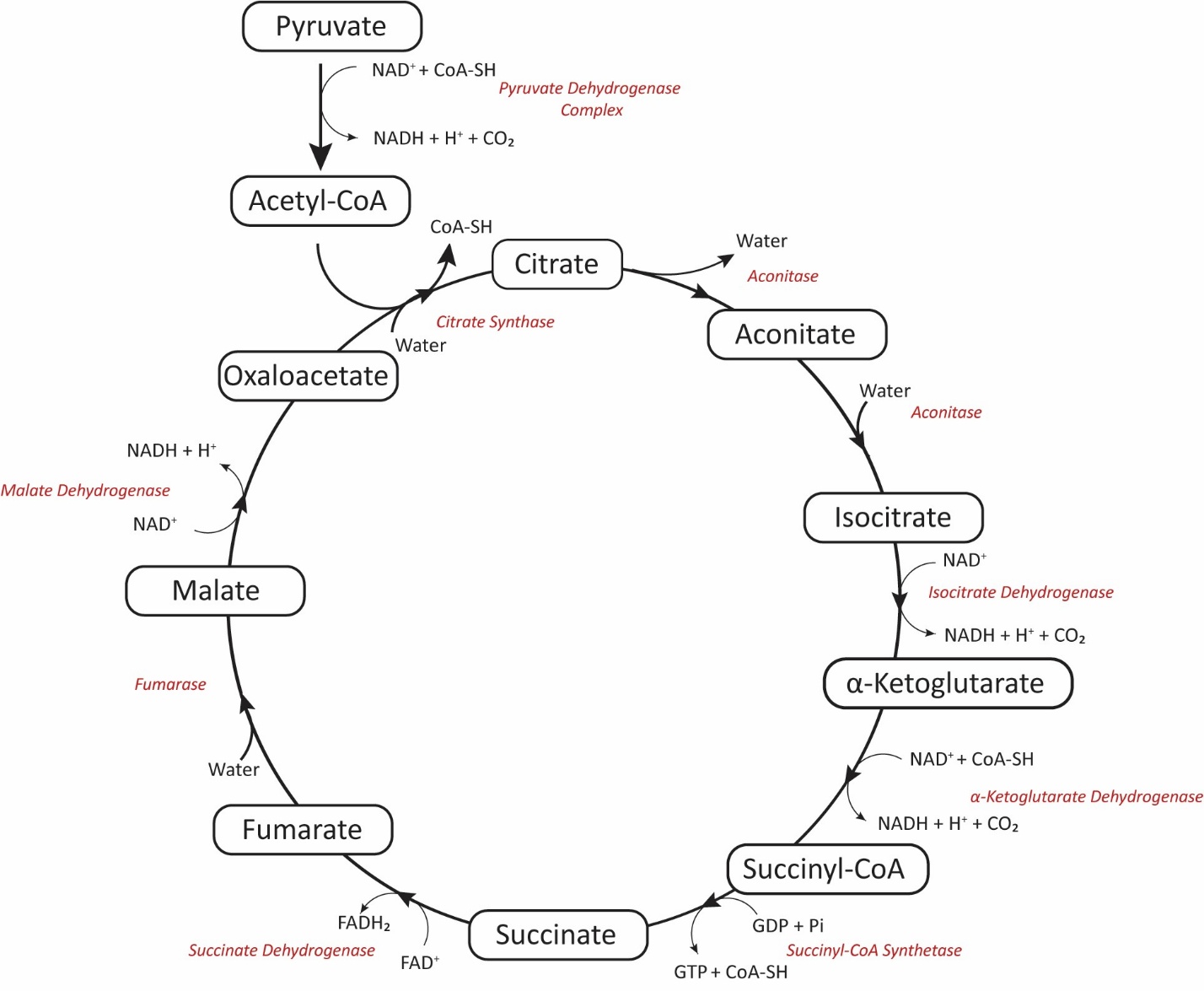

Supplemental Figure 1 – Krebs cycle schematic

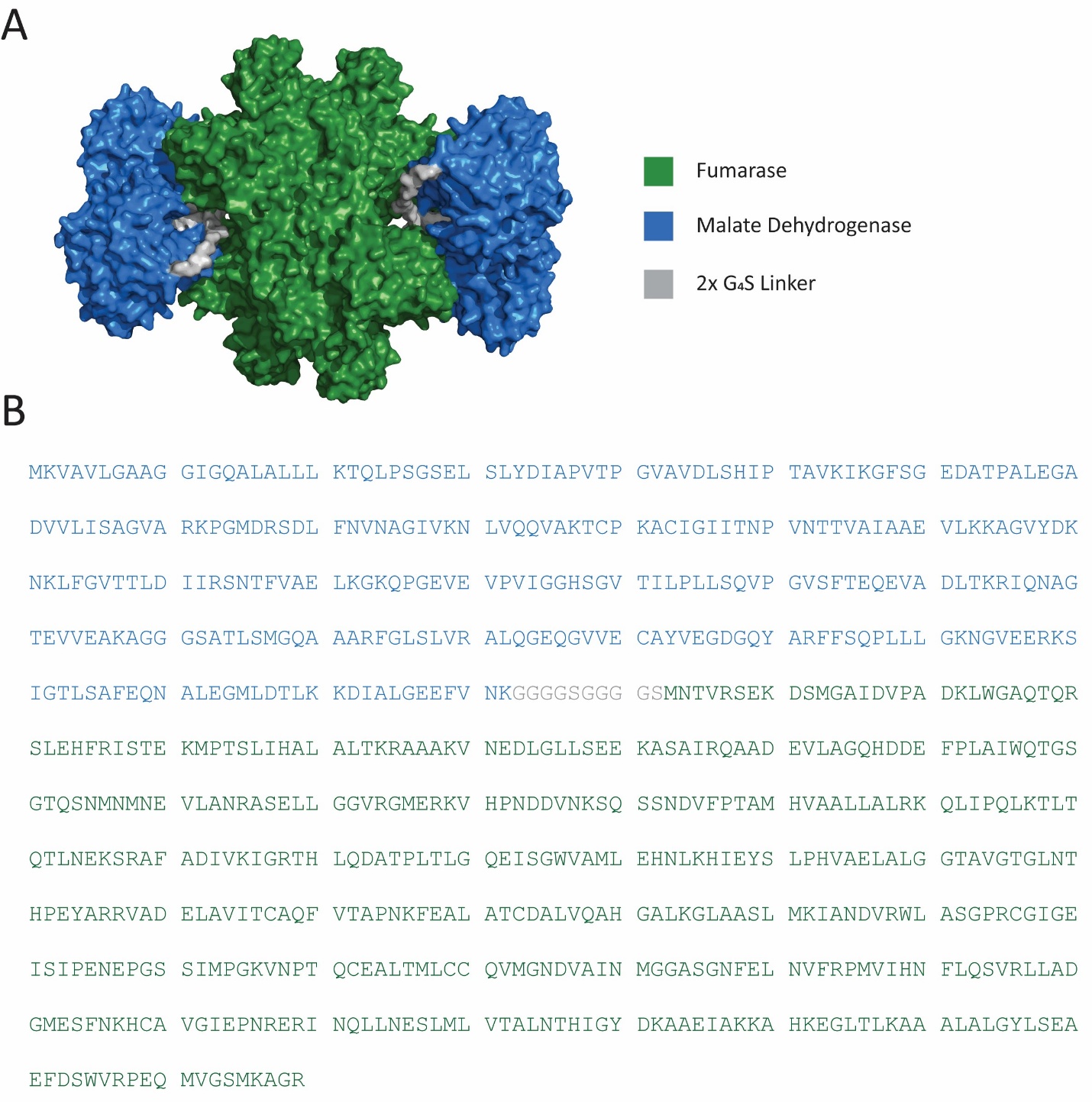

Supplemental Figure 2 – **(A)** Alphafold3 structure prediction of V0 **(B)** Protein sequence of the V0 construct

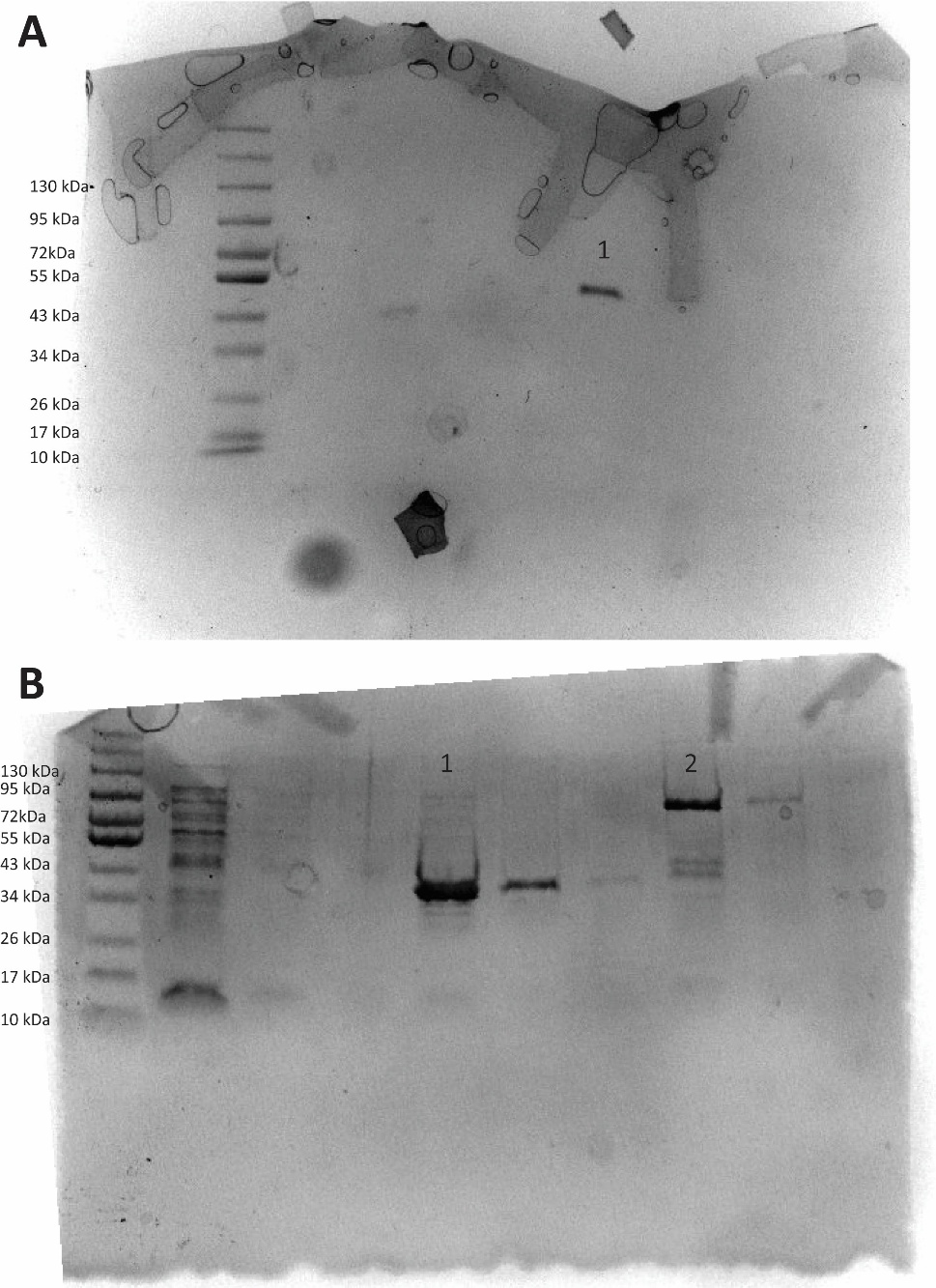

Supplemental Figure 3 – (**A**) Lane 1 - WT tetrameric Fumarase from E. coli. Expected MW 51 kDa (**B**) Lane 1 - Malate dehydrogenase (MDH), expected molecular weight 36 kDa. Lane 2 – V0 fusion. Expected molecular weight 90 kDa

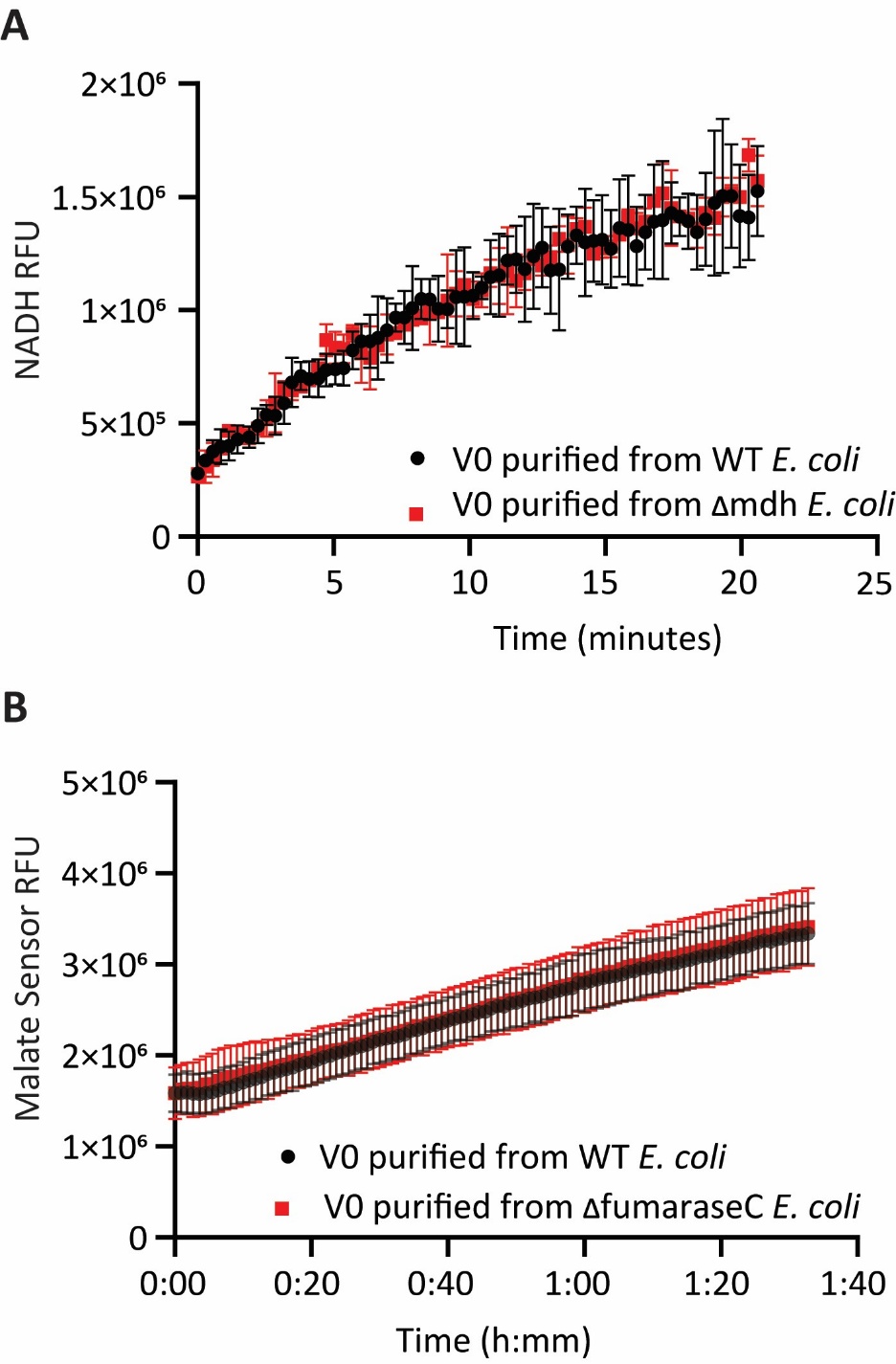

Supplemental Figure 4 – (**A**) V0 MDH turnover of malate and NAD+ to oxaloacetate and NADH. V0 was either purified from wild-type E. coli or a strain of E. coli lacking MDH (**B**) V0 fumarase turnover of fumarate to malate. V0 was either purified from wild-type E. coli or a strain of E. coli lacking FumC

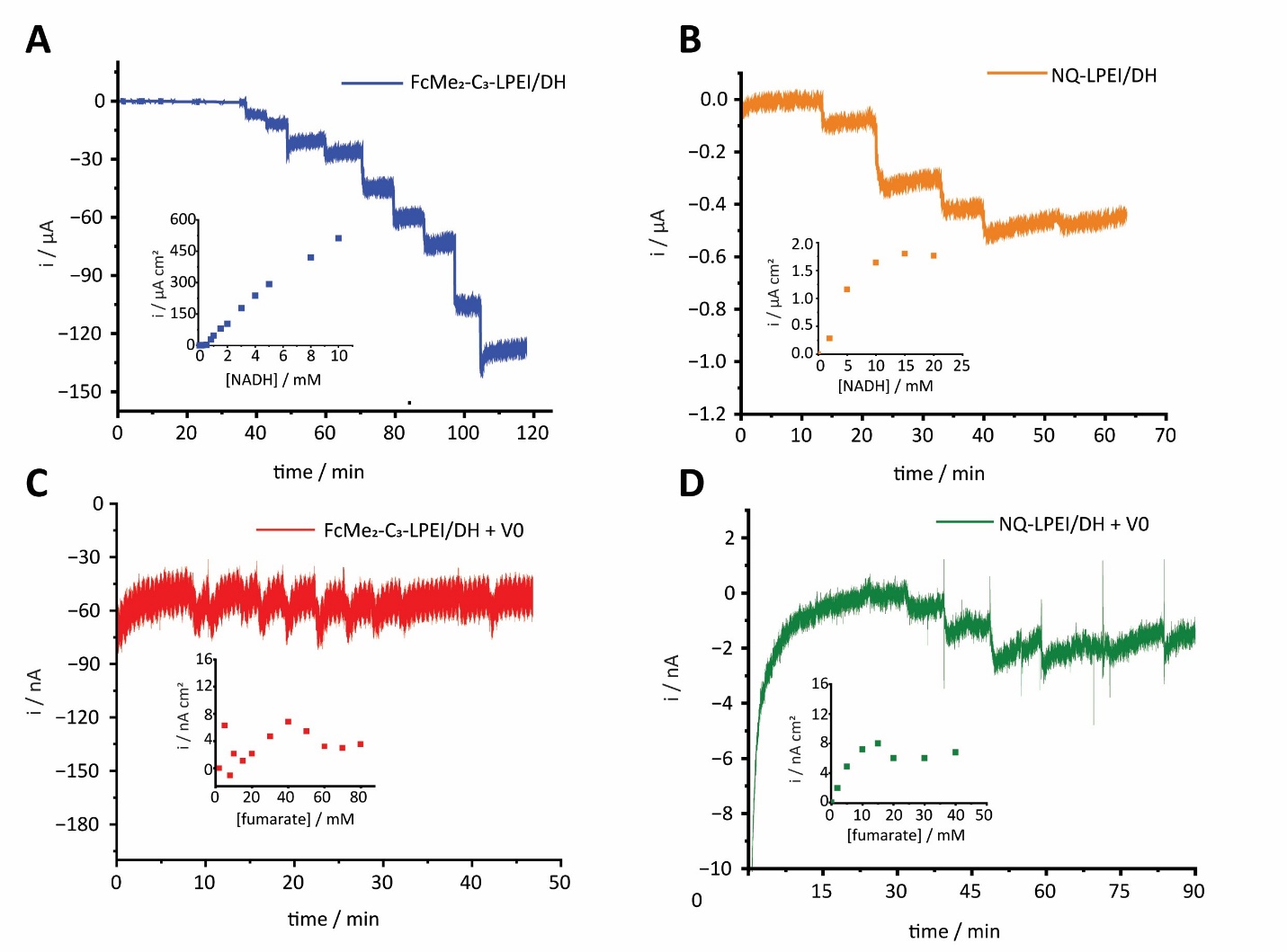

Supplemental Figure 5 - Amperometric j-t curves and corresponding calibration curves obtained upon increasing substrate concentrations (+0.25 vs SCE). Electrochemical evaluation of NADH oxidation with (**A**) FcMe_2_-C_3_-LPEI and (**B**) NQ-LPEI-LPEI. Both with diaphorase cross-linked. Electrochemical evaluation of fumarate oxidation and the same polymers with V0 crosslinked (**C**) FcMe_2_-C_3_-LPEI + V0 (**D**) NQ-LPEI + V0.

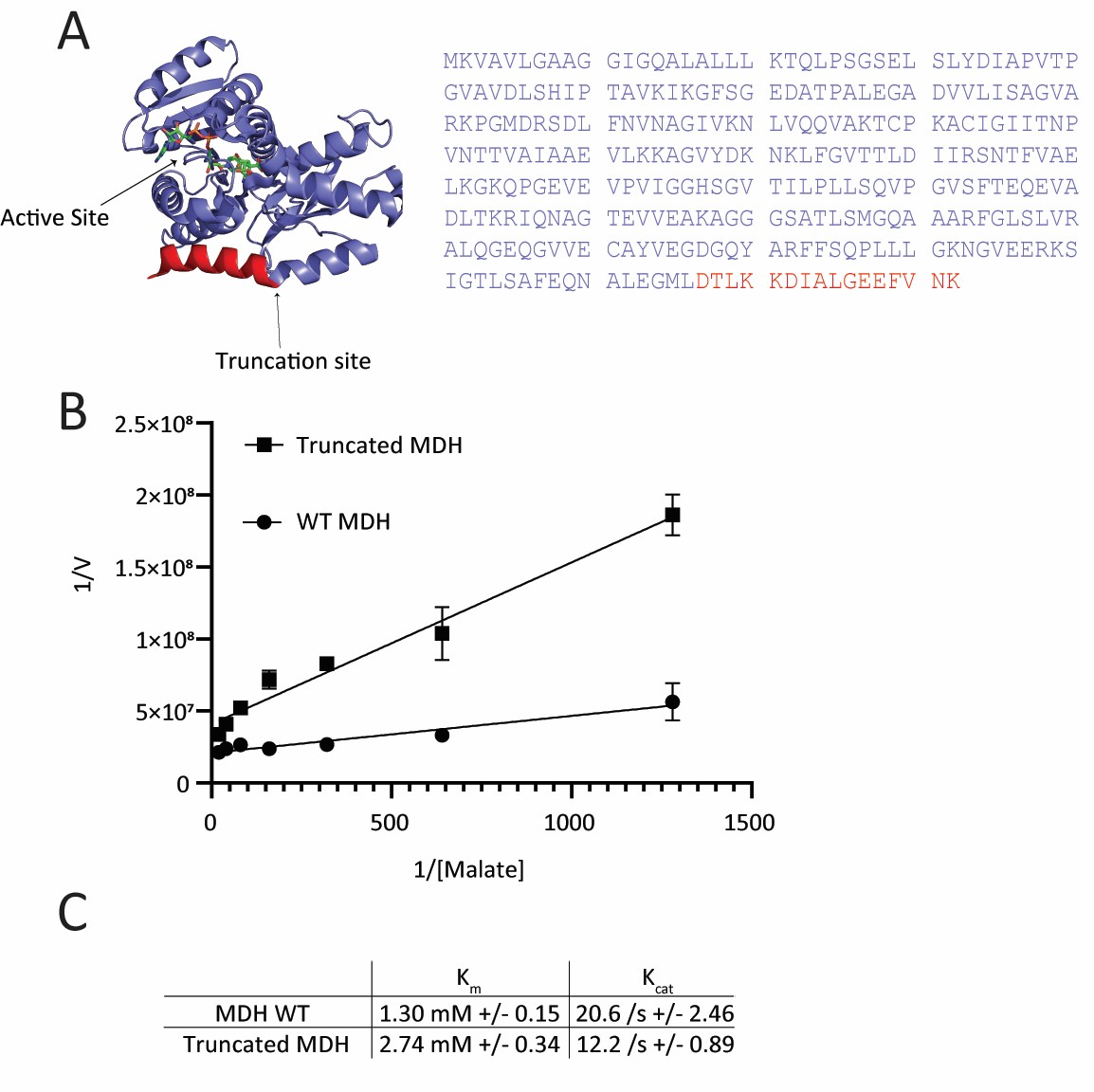

Supplemental Figure 6 - **(A)** Crystal structure of malate dehydrogenase (1EMD) labeled with targeted truncation site in red and **(B-C)** the effect on enzymatic kinetics

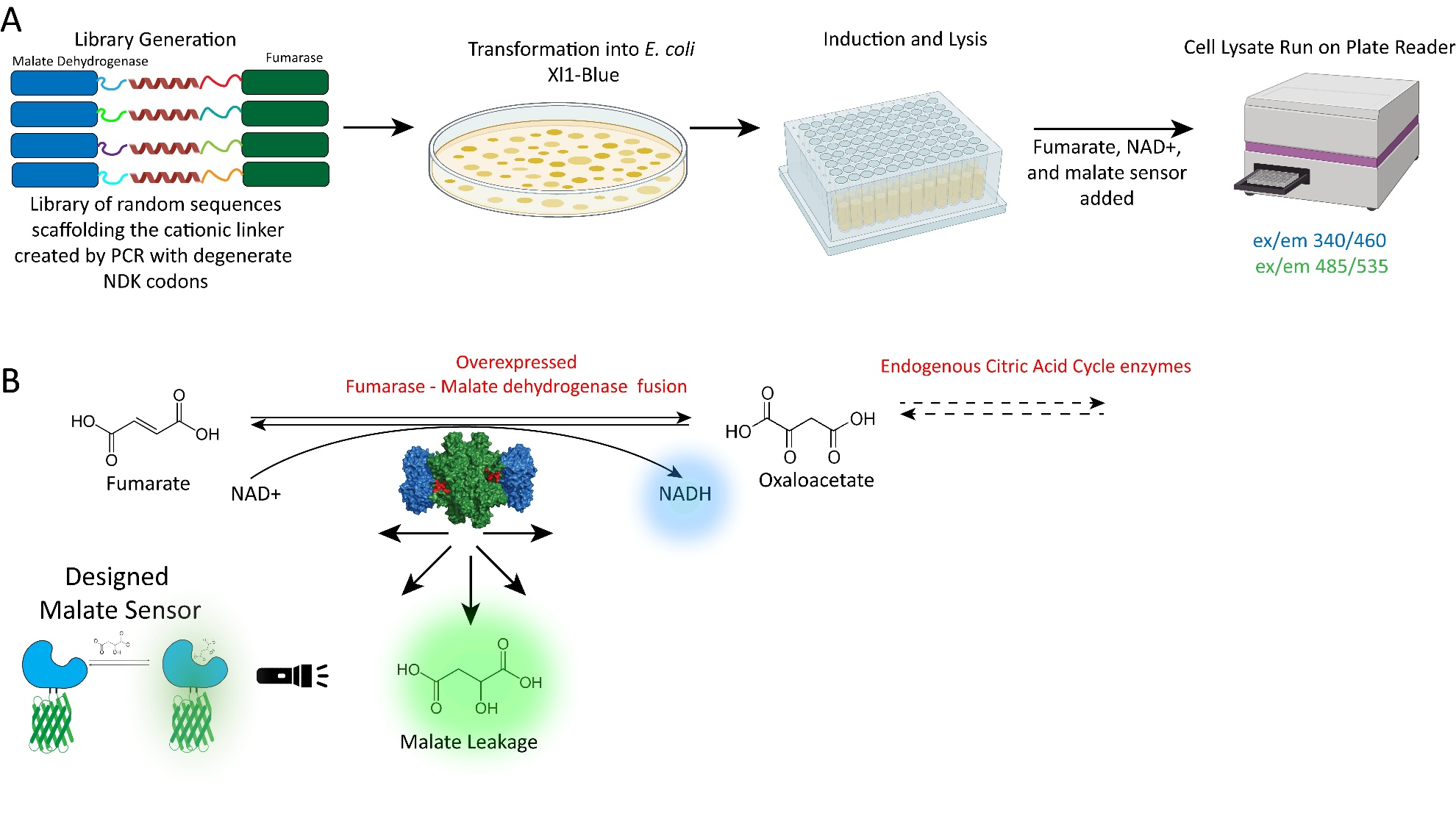

Supplemental Figure 7 – General workflow library generation and substrate channeling screening **(A)** A library of fusions between MDH and fumarase was generated by PCR with random scaffolding sequences. This library was then transformed into E. coli cells. Individual colonies were picked and grown in plates inducing the fusion protein production. Cultures were lysed, and added to plates containing NAD+, Fumarate, and a fluorescent malate sensor. The plates were run on a plate reader measuring 340/460 ex/em to quantify NADH production, as well as the 500/540 ex/em to measure the malate leakage from the fusion enzymes by the malate biosensor. **(B)** Reaction scheme of the conversion of fumarate to malate and oxaloacetate by the fumarase malate dehydrogenase fusion. Malate and NADH are highlighted with the corresponding fluorescence colors used to detect their formation

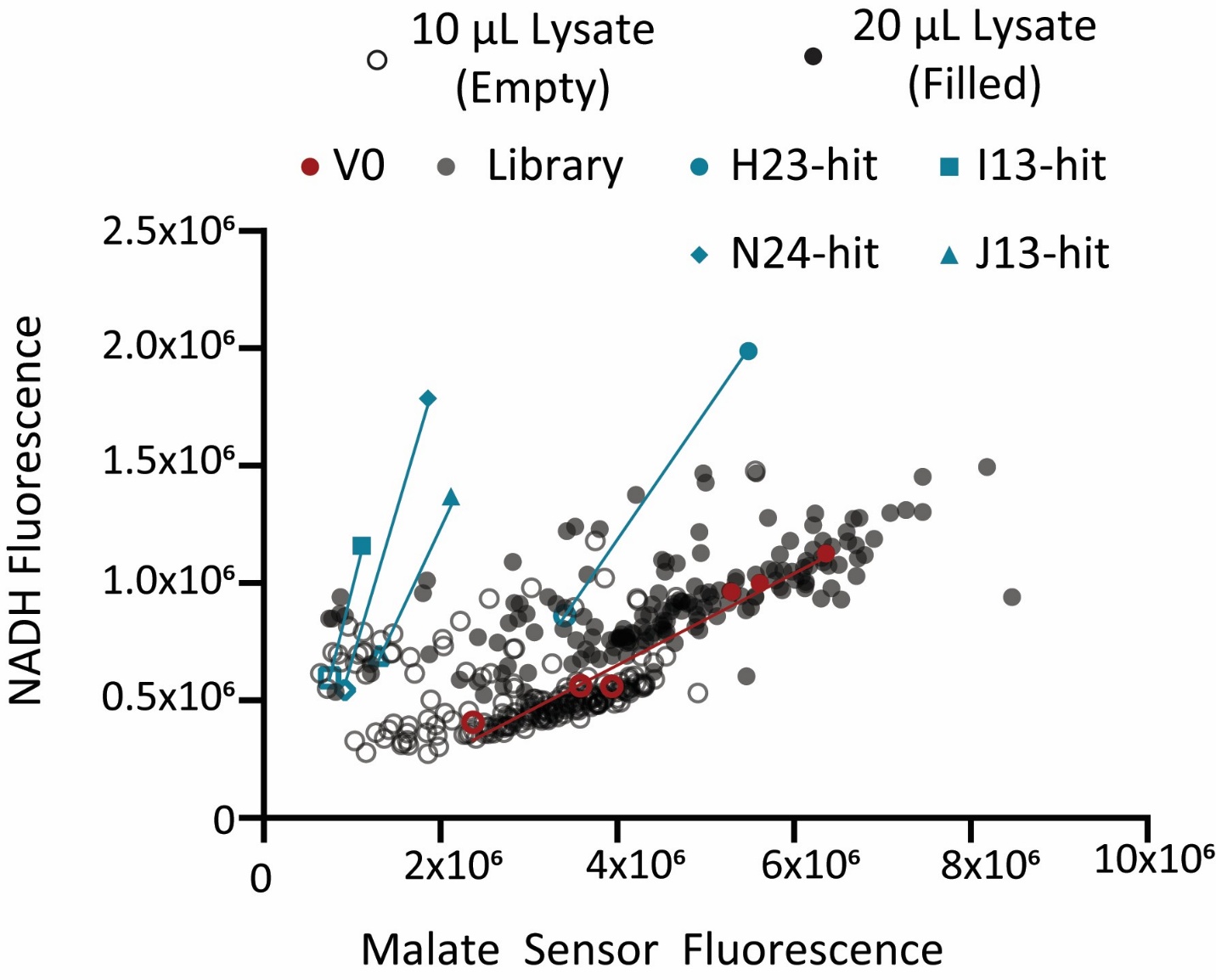

Supplemental Figure 8 – repeat run of figure 2C

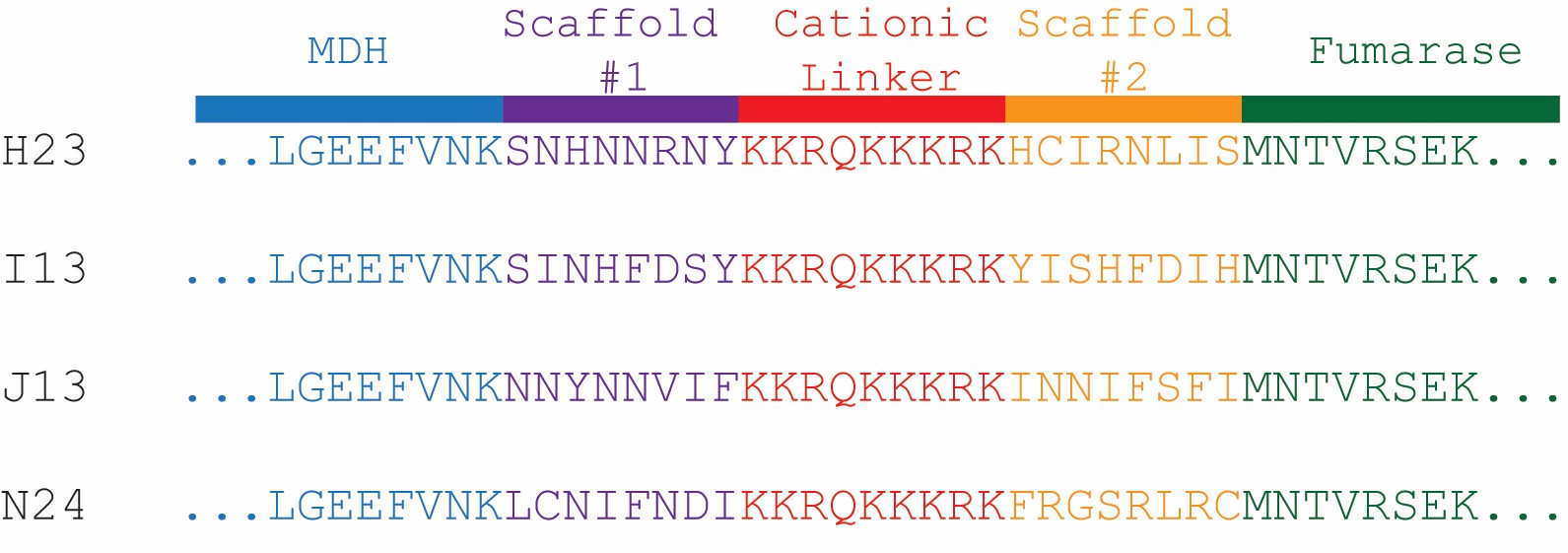

Supplemental Figure 9 – Scaffold sequences from hits identified through lysate screen

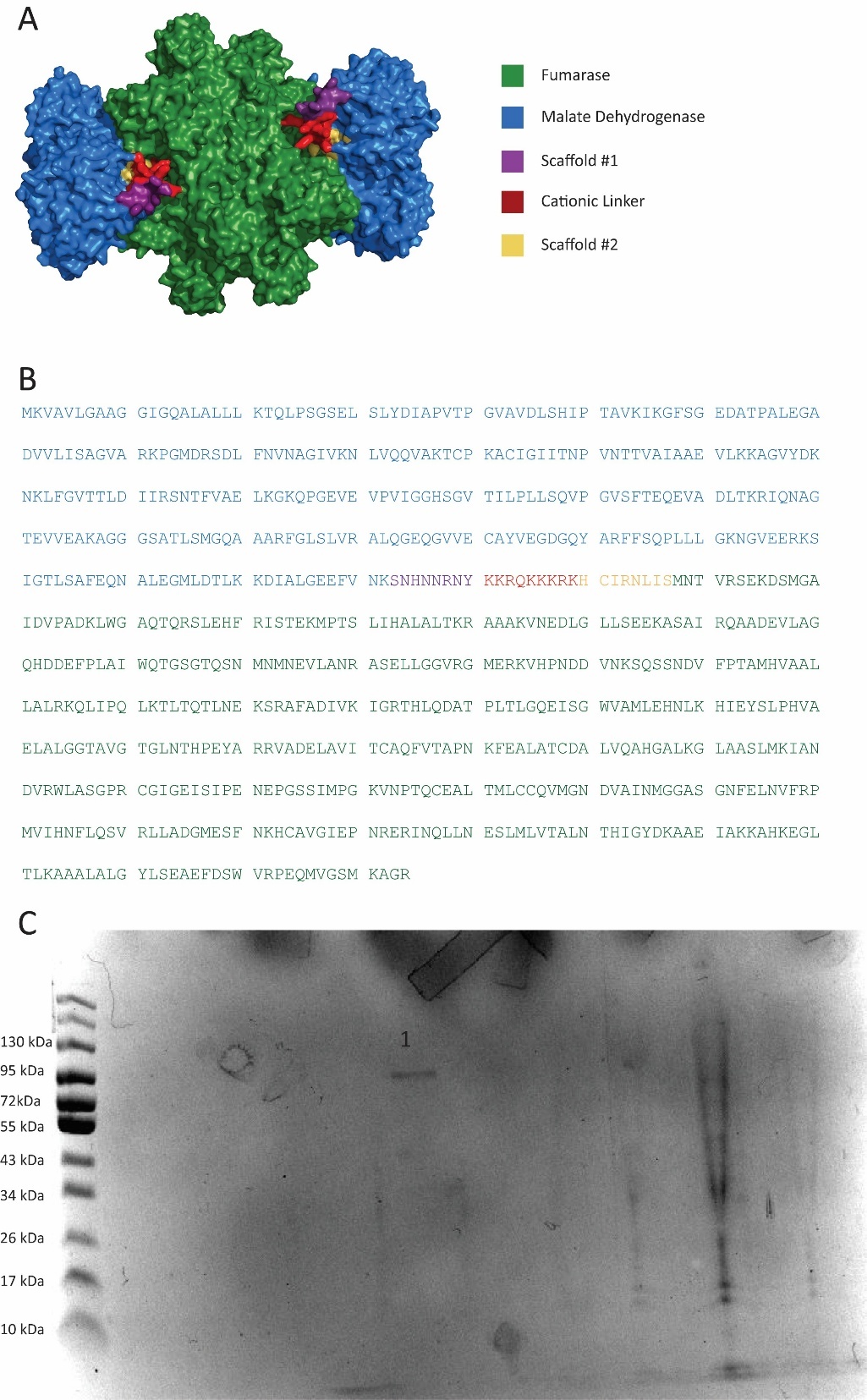

Supplemental Figure 10 – **(A)** Alphafold3 structure H23 construct **(B)** Sequence of the H23 construct (**C**) Gel purification of H23. Lane 1 – H23 fusion protein. Expected molecular weight 90 kDa

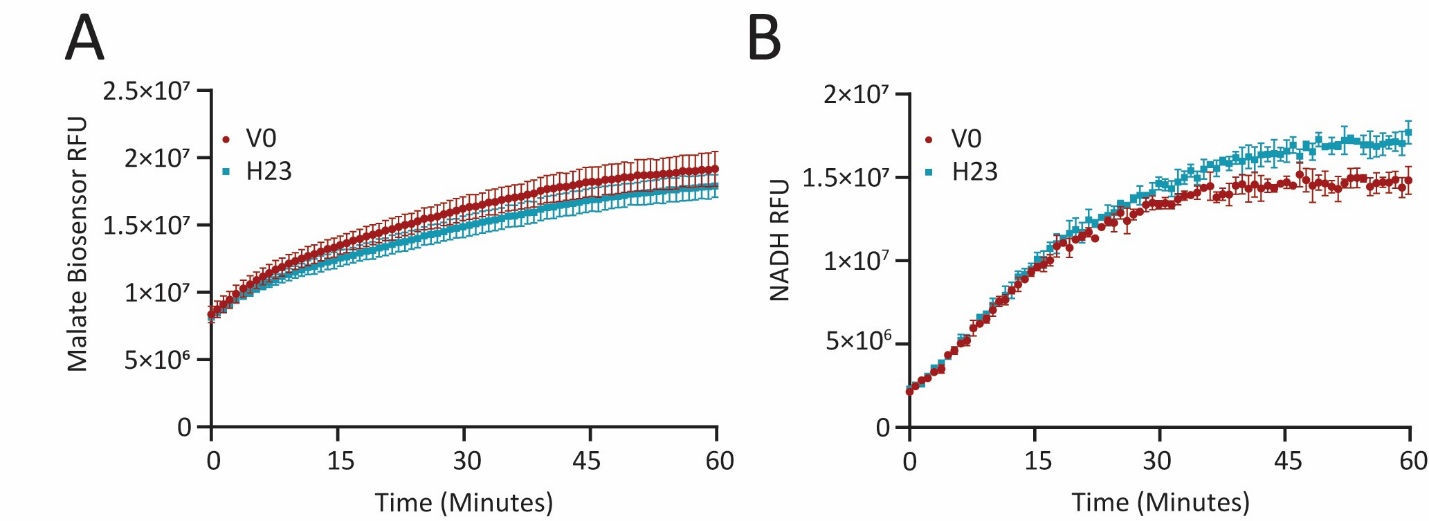

Supplemental Figure 11 – Comparison of **(A)** malate and **(B)** NADH production for both V0 and H23 from the addition of fumarate and NAD+

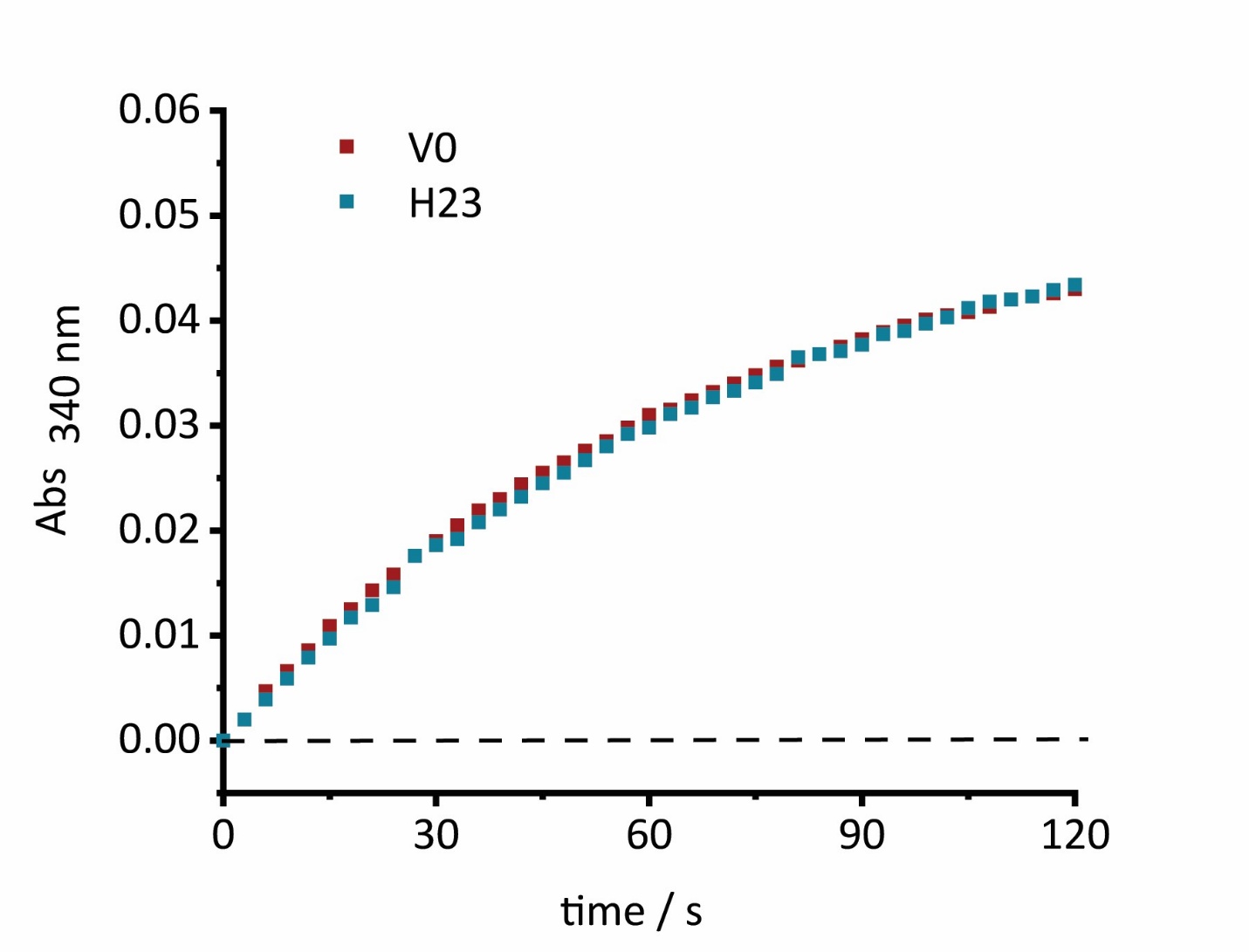

Supplementary Figure 12 – Transient time analysis of V0 and H23 with malate injection

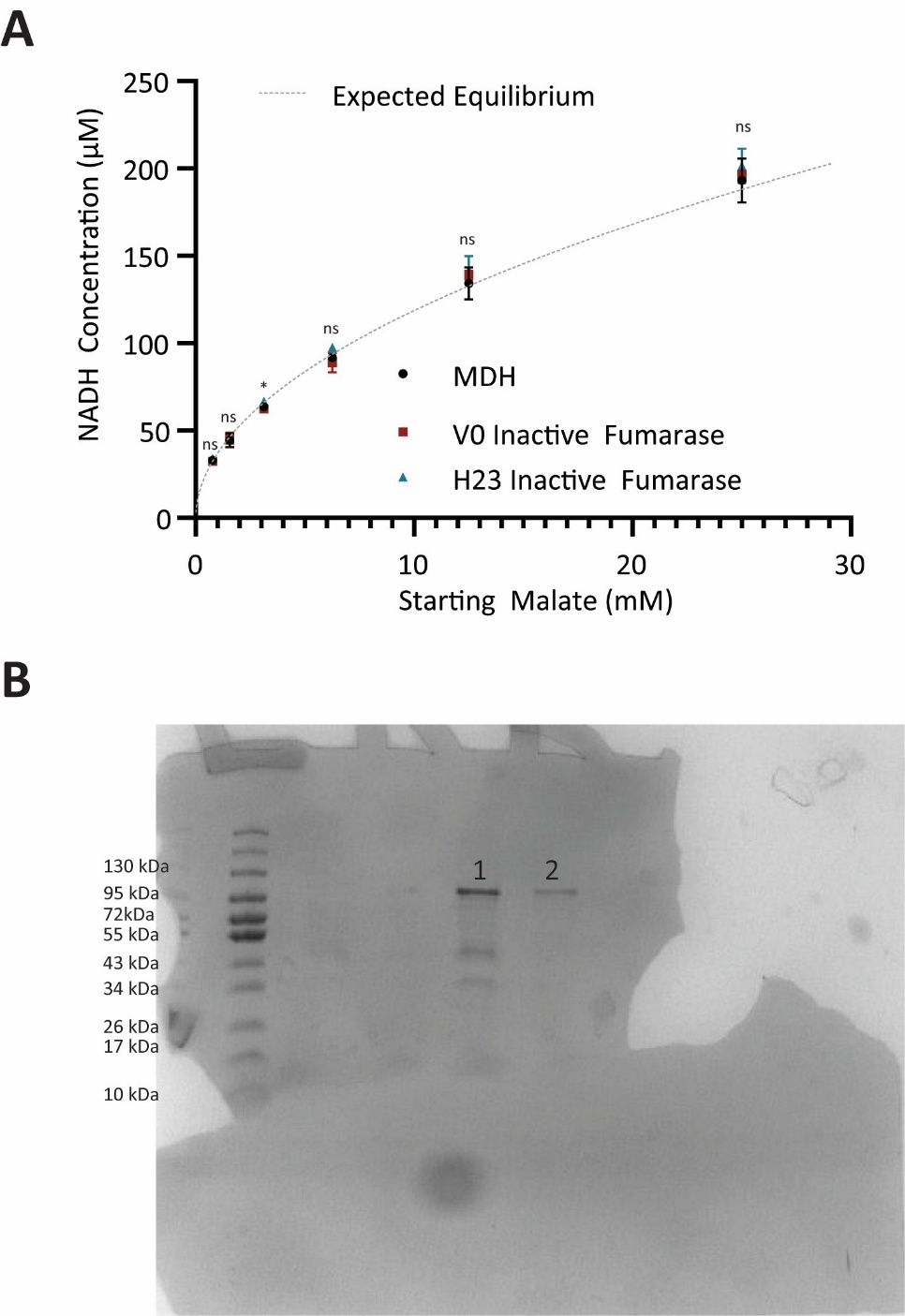

Supplemental Figure 13 – (**A**) Comparison of equilibrium NADH concentrations after enzymatic reaction with malate and NAD. V0 and H23 had inactivating mutations in the active site of their fumarase. Statistical significance of H23 vs MDH denoted above each point (**B**) Gel verification of inactivating mutants in fumarase. Four mutations were performed in fumarase to ensure complete inactivation: S318A, S319A, K324A, and N326A. Lane 1 – H23 mutant - Expected Molecular Weight 90 kDa. Lane 2 – V0 mutant - Expected Molecular Weight 90 kDa

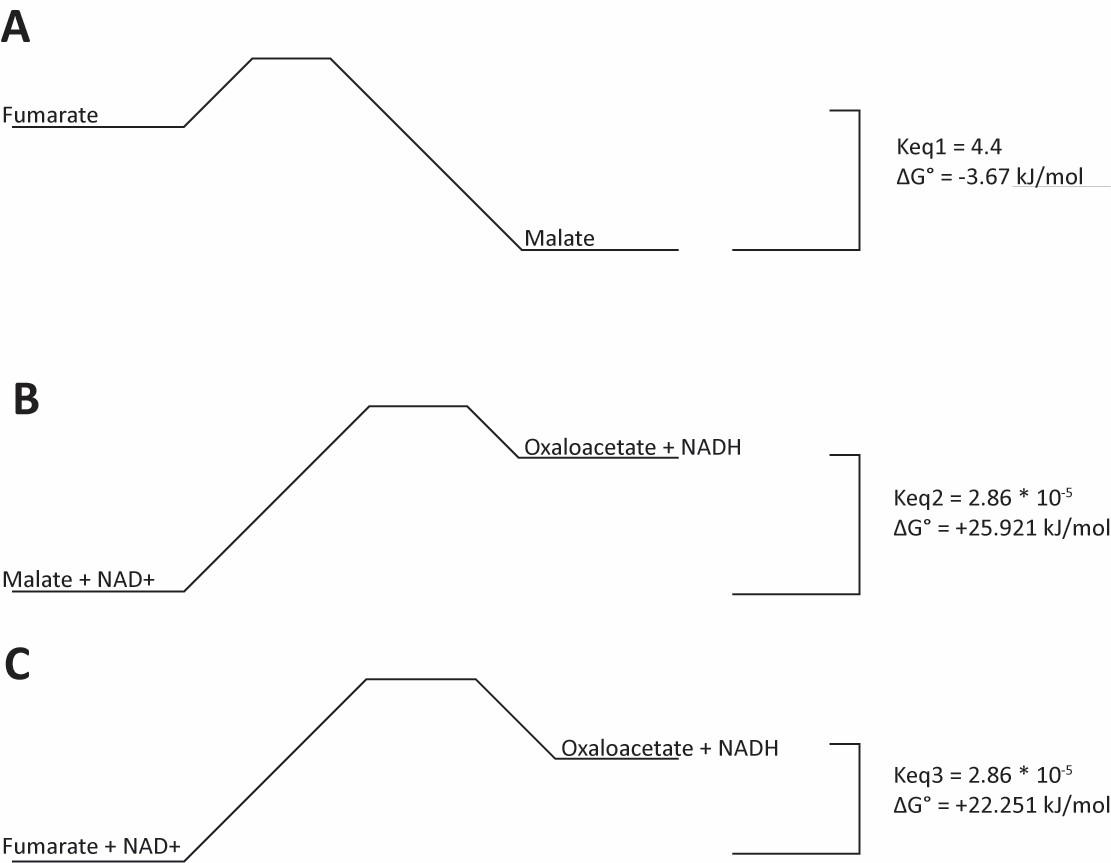

Supplemental Figure 14 – ΔG° and Keq calculated for the conversion of different reactions **(A)** Known Keq for the conversion of fumarate to malate was used to calculate ΔG° **(B)** Known Keq for the conversion of malate and NAD^+^ to oxaloacetate and NADH was used to calculated ΔG° **(C)** In a perfectly channeling system, for the conversion of fumarate and NAD+ to oxaloacetate and NADH, the ΔG° would be the sum of ΔG° from **(A)** and **(B)**. Keq for a perfectly channeled system can be calculated from the predicted ΔG°.

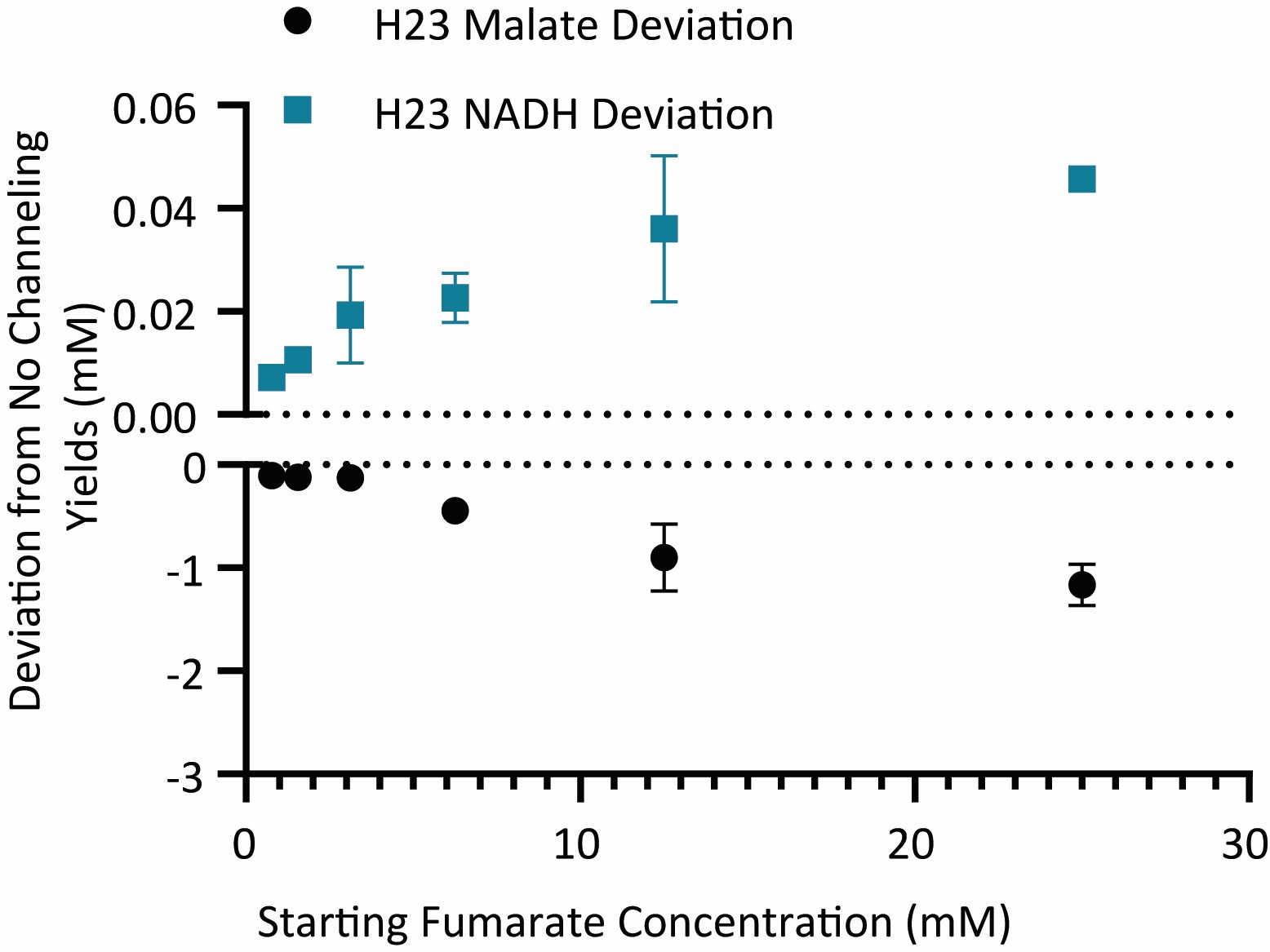

Supplemental Figure 15 - Deviation of H23 from the expected yields for a system with no channeling for both malate and NADH

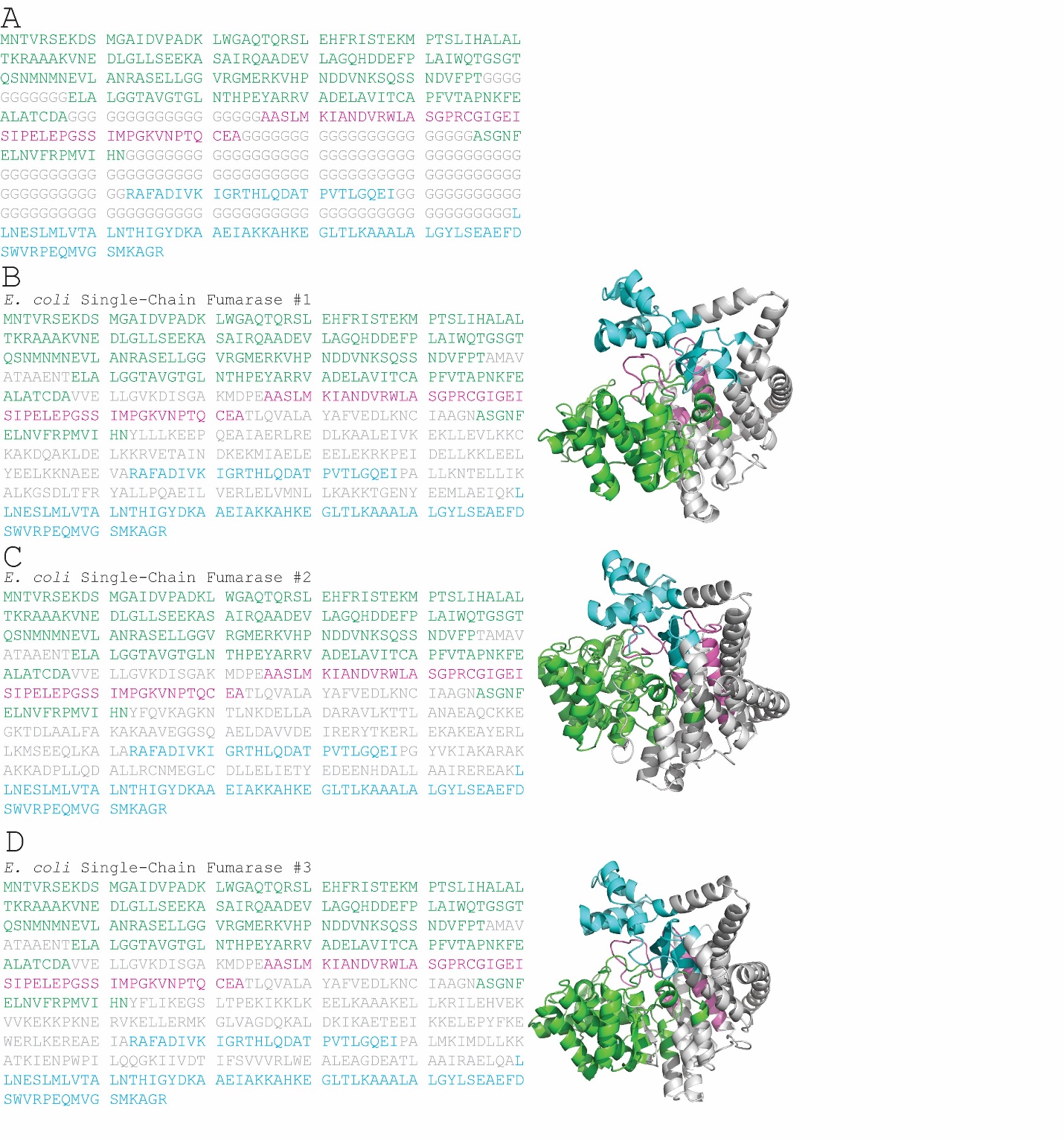

Supplemental Figure 16 – Design of E. coli single-chain fumarase. Fragments originating from subunit 1 of the tetrameric fumarase are colored in green, fragments from subunit 2 are colored in blue, and fragments from subunit 3 are colored in pink. Residues that were de novo designed are colored in grey (**A**) Base sequence and fragments used for designing E. coli single-chain fumarase. Greyed-out glycine indicates spatial placeholders designed by RFDiffusion. (**B-D**) Sequences of single-chain E. coli fumarase (**B**) #1 (**C**) #2 (**D**) and #3. Glycine placeholders were replaced with optimized residues predicted to stabilize the designed fold by ProteinMPNN, in addition to predicted Alphafold2 structures.

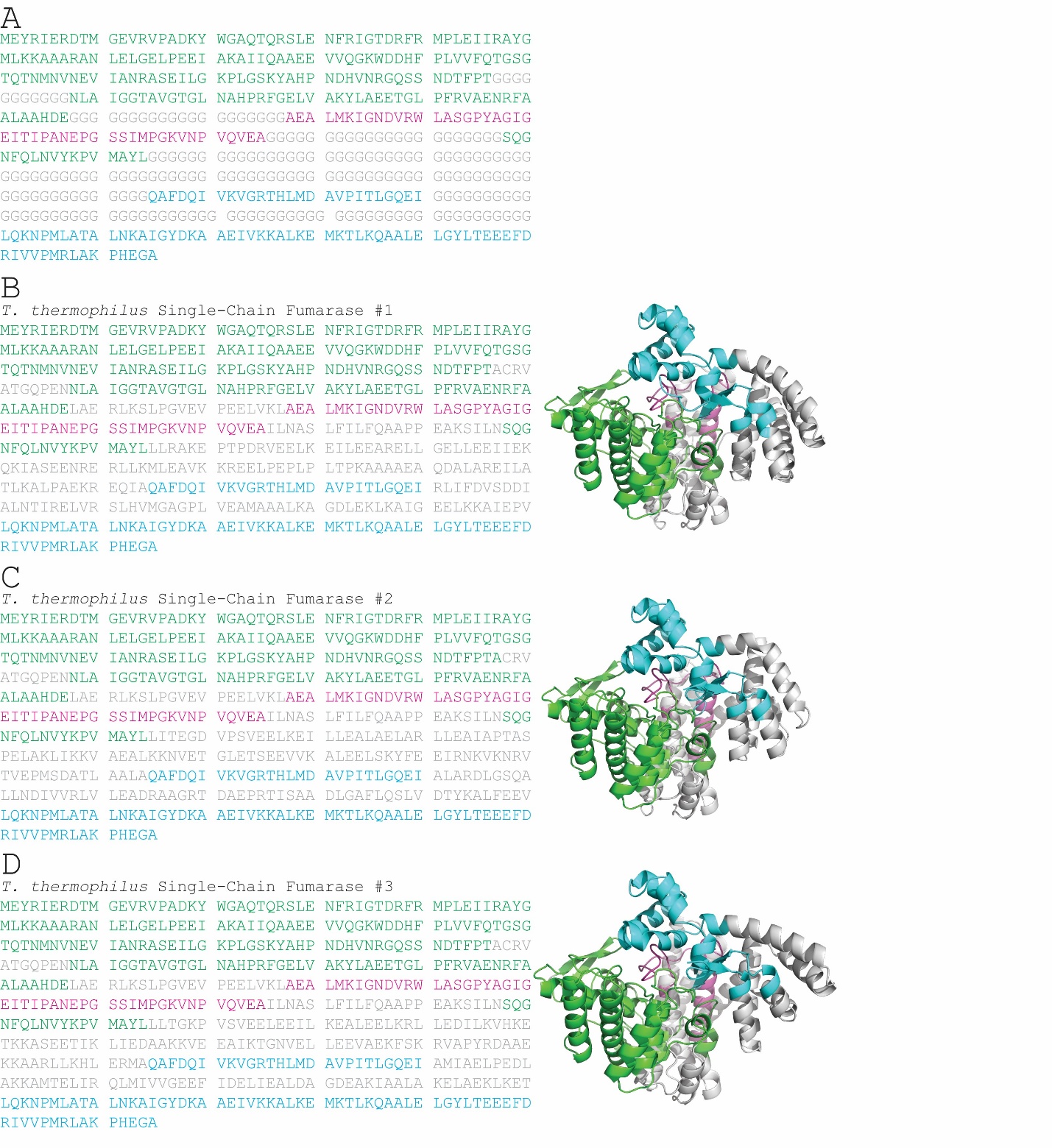

Supplemental Figure 17 - Design of T. thermophilus single-chain fumarase. Fragments originating from subunit 1 of the tetrameric fumarase are colored in green, fragments from subunit 2 are colored in blue, and fragments from subunit 3 are colored in pink. Residues that were de novo designed are colored in grey (**A**) Base sequence and fragments used for designing T. thermophilus single-chain fumarase. Greyed out glycine indicates spatial placeholders designed by RFDiffusion. (**B-D**) Sequences of single-chain T. thermophilus fumarase (**B**) #1 (**C**) #2 (**D**) and #3. Glycine placeholders were replaced with optimized residues predicted to stabilize the designed fold by ProteinMPNN, in addition to predicted Alphafold2 structures.

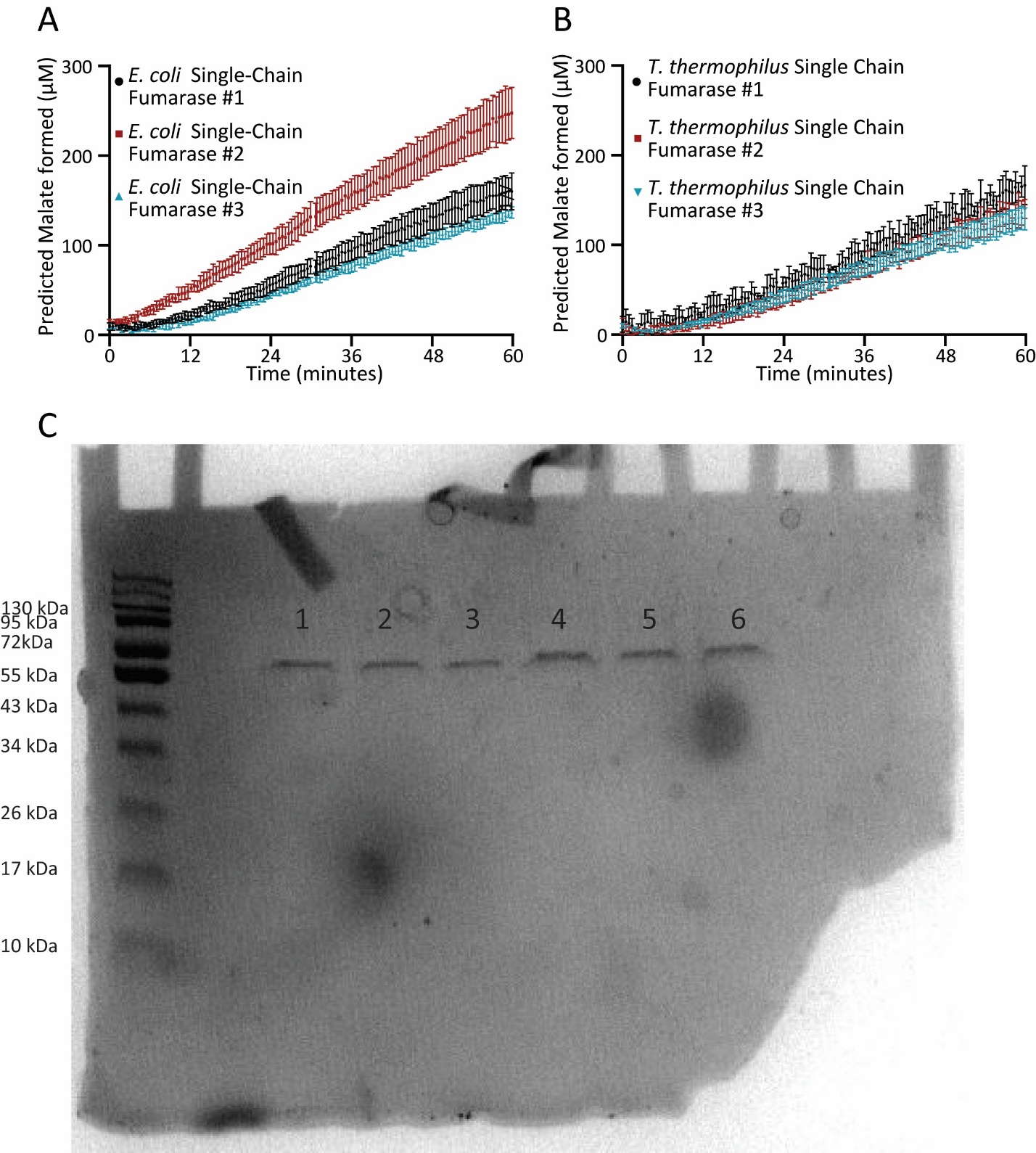

Supplemental Figure 18 – Initial activity screen of (**A**) E. coli based single-chain fumarase(**B**) T. thermophilus based single-chain fumarase. (**C**) Gel verification of single-chain fumarase to ensure purity.

Lane1 – Thermophilus single-chain #1 Expected Molecular Weight 62.8 kDa

Lane 2 – Thermo Thermophilus single-chain #2 Expected Molecular Weight 62.4 kDa

Lane 3 – Thermo Thermophilus single-chain #3 Expected Molecular Weight 62.5

Lane 4 - E. coli single-chain #1 Expected Molecular Weight 62.8 kDa

Lane 5 - E. coli single-chain #2 Expected Molecular Weight 61.9 kDa

Lane 6 - E. coli single-chain #3 Expected Molecular Weight 61.8 kDa

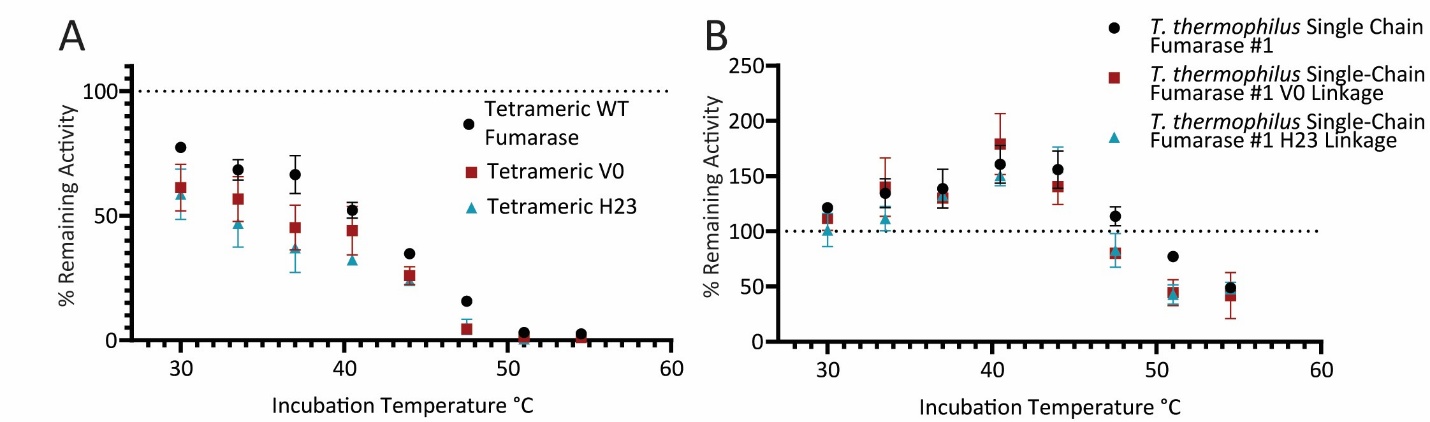

Supplemental Figure 19 – Relative activity of fumarase was compared after heat incubation. Unlinked fumarases were compared to fumarases linked to MDH with both the V0 and H23 linkers (**A**) tetrameric fumarase (**B**) Single-chain T. thermophilus fumarase

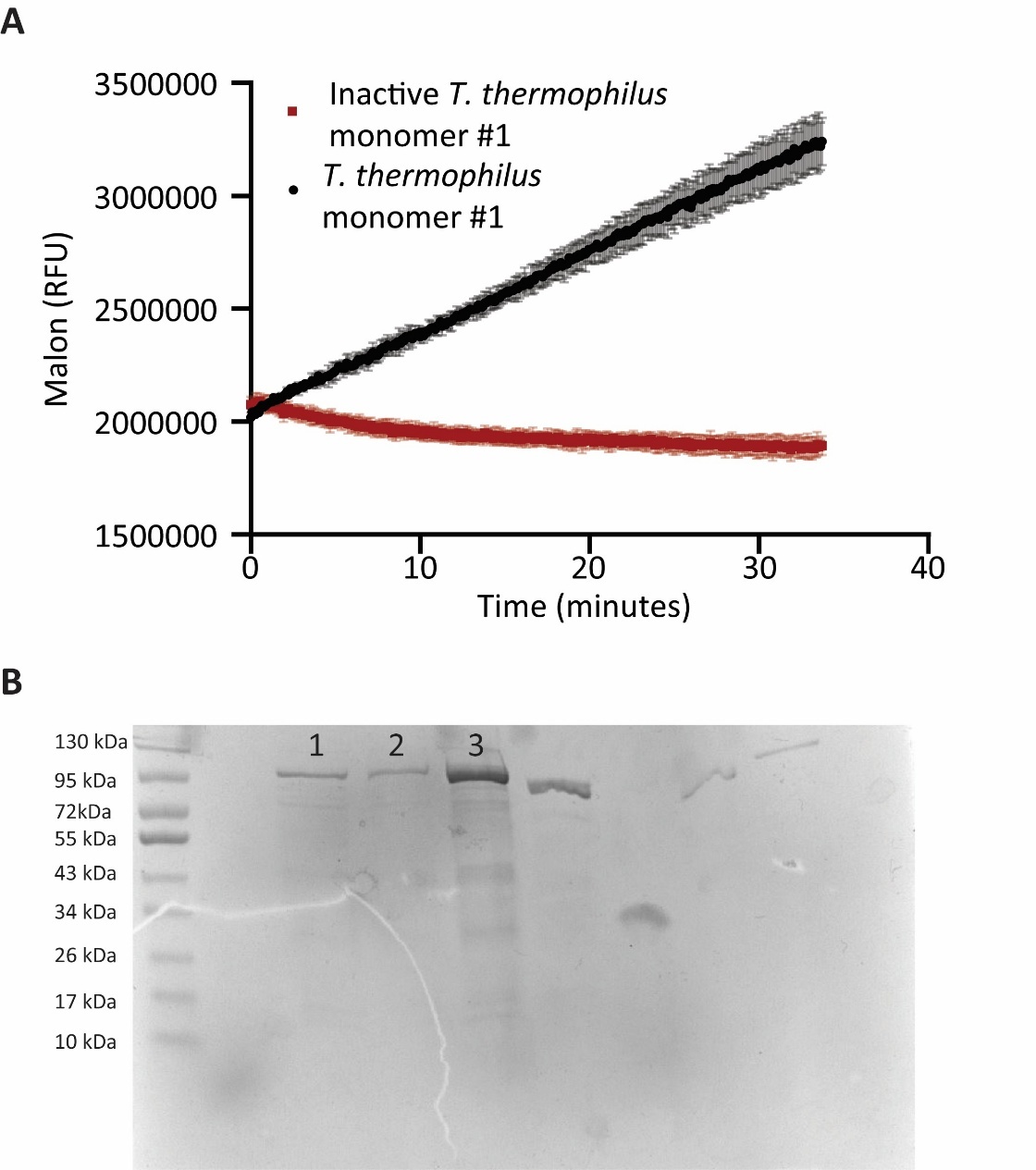

Supplemental Figure 20 (**A**) Comparison of T. thermophilus single-chain fumarsae #1 to a variant with inactivating mutations. Both versions were connected to MDH with the V0 linker (**B**) Gel verification of T. Thermophilus single-chain #1 Linked to MDH using V0 linkage (Lane 1), H23 linkage (Lane 2) and V0 linkage but with inactivating mutations in the active site (Lane 3). Inactivating mutations correspond to mutations previously used in tetrameric fumarase. Using single-chain numbering, 4 mutations were made S261A, S262A, K267A, N269A to ensure complete inactivation

|  | Yield Increase |
| --- | --- |
| H23 – in Solution | 29% +/- 8% |
| *T. thermophilus* single-chain #1 H23 – in Solution | 63% +/- 25 % |
| Theoretical Perfect Channeling | 130% |

*Supplemental Table 1 – Enhanced yield (current or NADH) of tetrameric and* single-chain *fumarase on an electrode and in solution compared to non-channeling controls. Additionally, theoretical perfect channeling yield increases were calculated (see methods)*

|  | RMSD Alignment Å | |
| --- | --- | --- |
|  | Approximate tetramer used in design | PDB:4APB |
| *E. coli* single-chain #1 | 0.412 | 0.841 +733 +/- 0.033 |
| *E. coli* single-chain #2 | 0.418 | 0.776 +/- 0.036 |
| *E. coli* single-chain #3 | 0.386 | 0.792 +/- 0.039 |
| *T. thermophilus* single-chain #1 | 0.361 | 0.755 +/- 0.089 |
| *T. thermophilus* single-chain #2 | 0.39 | 0.752 +/- 0.142 |
| *T. thermophilus* single-chain #3 | 0.428 | 0.733 +/- 0.111 |

Supplemental Table 2 – *Root-mean-square deviation (RMSD) values from structural alignments of AlphaFold2-predicted* single-chain *fumarase models to (1) the approximate tetrameric models of E. coli and T. thermophilus fumarases (constructed by aligning PDB 1YFE and 1VDK subunits to PDB 4APB), and (2) the crystal structure of M. tuberculosis fumarase (PDB 4APB). Standard errors for the second column were derived by aligning monomeric models to each of the four pockets in 4APB.*

| **Primer Name** | **Sequence** | **Purpose** |
| --- | --- | --- |
| NJR_167a | CAGTGCTGCAATGATACCGC | Backbone Primer - pairs with multiple primers in this study |
| NJR_168a | AGACTGGATGGAGGCGGATA | Backbone Primer - pairs with multiple primers in this study |
| NJR_257 | TAAGAAGGAGATATGGATCCATGAATACAGTACGCAGCGAAA | Amplification of FumC from E. coli genome pairs with NJR_258 |
| NJR_258 | GACCGTTTAAACTCAATGGTGATGGTGATGATGACGCCCGGCTTTCATACTG | Amplification of FumC from E. coli genome pairs with NJR_257 |
| NJR_259 | ACTTTAAGAAGGAGATATGGATCCATGGGGGCGATTGATGTCC | Amplification of FumC with 10 amino acid truncation, pairs with NJR_258 |
| NJR_255 | TTGAGTTTAAACGGTCTCCAGC | pBad Backbone amplification, pairs with NJR_256 |
| NJR_256 | TGGATCCATATCTCCTTCTTAAAGT | pBad Backbone amplification, pairs with NJR_255 |
| NJR_262 | TGACGATAAGGATCCGAGCTCGAGAATGAAAGTCGCAGTCCTCGG | Amplification of MDH from E. coli genome pairs with NJR_263 |
| NJR_263 | TCATCCGCCAAAACAGCCAAGCTTTTACTTATTAACGAACTCTTCGCCCA | Amplification of MDH from E. coli genome pairs with NJR_262 |
| NJR_264 | GCCAAAACAGCCAAGCTTTTACAGCATACCTTCCAGCGCGTTCT | Amplification of MDH with 10 amino acid truncation, pairs with NJR_262 |
| NJR_260 | TAAAAGCTTGGCTGTTTTGGCG | pBad Backbone amplification, pairs with NJR_260 |
| NJR_261 | TCTCGAGCTCGGATCCTTATC | pBad Backbone amplification, pairs with NJR_261 |
| NJR_273 | GGAACCGCCCCCACCACTGCCACCACCTCCCTTATTAACGAACTCTTCGCCCA | Fusion of MDH and Fumarase pairs with NJR_168a |
| NJR_274 | GGAGGTGGTGGCAGTGGTGGGGGCGGTTCCATGAATACAGTACGCAGCGA | Fusion of MDH and Fumarase pairs with NJR_167a |
| Primer 401a | AAGAAACGTCAGAAGAAAAAGCGTAAANDTNDTNDTNDTNDTNDTNDTNDTATGAATACAGTACGCAGCGAAAAAGA | Amplification of Randomized Linker 1 - pairs with NJR_167a |
| Primer 402a | TTTACGCTTTTTCTTCTGACGTTTCTTAHNAHNAHNAHNAHNAHNAHNAHNCTTATTAACGAACTCTTCGCCCAGG | Amplification of Randomized Linker 2 - pairs with NJR_168a |
| NJR_439 | GAACTGGAGCCGGGTGCGGCGATTATGCCGGGCAAAGTTAACCCCA | Removes fumarase active site (E. coli variants) pairs with NJR_167a |
| NJR_440 | GTTAACTTTGCCCGGCATAATCGCCGCACCCGGCTCCAGTTCAGGAATA | Removes fumarase active site (E. coli variants) pairs with NJR_168a |
| NJR_445 | TGAACCGGGTGCAGCGATCATGCCAGGTAAGGTTAATCCCG | Removes fumarase monomer activity (Thermo variants) pairs with NJR_167a |
| NJR_446 | CCTGGCATGATCGCTGCACCCGGTTCATTGGCAGGGATGGT | Removes fumarase monomer activity (Thermo variants) pairs with NJR_168a |
| NJR_499 | ATGAATACAGTACGCAGCGA | Amplification of E. coli monomer pairs with NJR_542 |
| NJR_542 | TCTTCTCTCATCCGCCAAAACA | Amplification of E. coli monomer pairs with NJR_499 |
| NJR_543 | TAAACGGTCTCCAGCTTGGC | Amplification of MDH Backbone pairs with NJR_544a or NJR_544b |
| NJR_544a | CGCCCCCATCGAATCTTTTTCGCTGCGTACTGTATTCATGGAACCGCCCCCACCACTGCCACCACCTCCCTTAT | Amplification of MDH Backbone (use V0) pairs with NJR_514 |
| NJR_544b | TTTTTCGCTGCGTACTGTATTCATACTAATAAGATTACGAATACAATGTTTACGCTTTTTC | Amplification of MDH Backbone (use H23) pairs with NJR_543 |
| NJR_513 | ATGGAGTATCGTATCGAACGCGATACAATGGG | Amplification of T. thermophilus monomer pairs with NJR_514 |
| NJR_514 | TTTAAACTCATGCCCCCTCATGAGGCTTGGCTAAACGC | Amplification of T. thermophilus monomer pairs with NJR_513 |
| NJR_515 | CTCATGAGGGGGCATGAGTTTAAACGGTCTCCAGCTT | Amplification of MDH Backbone pairs with NJR_516 or NJR_518 |
| NJR_516 | ATCGCGTTCGATACGATACTCCATGGAACCGCCCCCACCACTGCCACCAC | Amplification of MDH Backbone (use V0) pairs with NJR_515 |
| NJR_518 \| CGCGTTCGATACGATACTCCATACTAATAAGATTACGAATACAATGTTTACGCTTTTTCTTCTGA | CGCGTTCGATACGATACTCCATACTAATAAGATTACGAATACAATGTTTACGCTTTTTCTTCTGA | Amplification of MDH Backbone (use H23) pairs with primer 515 |

Supplemental Table 3 – *Primers used in this study*

| **PCR Product 1** | | **PCR Product 2** | | **Product** |
| --- | --- | --- | --- | --- |
| **Forward** | **Reverse** | **Forward** | **Reverse** |  |
| NJR_257 | NJR_258 | NJR_255 | NJR_256 | pBAD Fumarase |
| NJR_259 | NJR_257 | NJR_255 | NJR_256 | pBad Truncated Fumarase |
| NJR_262 | NJR_263 | NJR_260 | NJR_261 | pBAD MDH |
| NJR_264 | NJR_263 | NJR_260 | NJR_261 | pBD Truncated MDH |
| NJR_273 | NJR_168a | NJR_274 | NJR_167a | pBAD V0 |
| NJR_401a | NJR_167a | NJR_402a | NJR_168a | pBAD Randon Scaffold Cationic Linker Library |
| NJR_439 | NJR_167a | NJR_440 | NJR_168a | pBAD Fumarase (*E. coli*) removed activity |
| NJR_445 | NJR_167a | NJR_446 | NJR_168a | pBAD Fumarase (*T. thermophilus*) removed activity |
| NJR_499 | NJR_542 | NJR_543 | NJR_544a | pBAD *E. coli* fumarase monomer V0 |
| NJR_499 | NJR_542 | NJR_543 | NJR_544b | pBAD *E. coli* fumarase monomer H23 |
| NJR_513 | NJR_514 | NJR_515 | NJR_516 | pBAD *T. thermophilus* fumarase monomer V0 |
| NJR_513 | NJR_514 | NJR_515 | NJR_518 | pBAD *T. thermophilus* fumarase monomer H23 |

Supplemental Table 4 – Gibson Assembly Table – *Products of various primers used together in Gibson assembly to form plasmid constructs used in this work*
